# A Myosin-7B dependent endocytosis pathway mediates cellular entry of α-Synuclein fibrils and polycation-bearing cargos

**DOI:** 10.1101/2020.02.12.946137

**Authors:** Qi Zhang, Yue Xu, Juhyung Lee, Michal Jarnik, Xufeng Wu, Juan S. Bonifacino, Jingshi Shen, Yihong Ye

## Abstract

Cell-to-cell transmission of misfolding-prone α-Synuclein (α-Syn) has emerged as a key pathological event in Parkinson’s disease. This process is initiated when α-Syn-bearing fibrils enter cells via clathrin-mediated endocytosis, but the underlying mechanisms are unclear. Using a CRISPR-mediated knockout screen, we identify SLC35B2 and Myosin-7B (MYO7B) as critical endocytosis regulators for α-Syn <u>p</u>re<u>f</u>ormed <u>f</u>ibrils (PFF). We show that SLC35B2, as a key regulator of heparan sulfate proteoglycan (HSPG) biosynthesis, is essential for recruiting α-Syn PFF to the cell surface because this process is mediated by interactions between negatively charged sugar moieties of HSPGs and clustered K-T-K motifs in α-Syn PFF. By contrast, MYO7B regulates α-Syn PFF entry by maintaining a plasma-membrane-associated actin network that controls membrane dynamics. Without MYO7B or actin filaments, many clathrin-coated pits fail to be severed from the membrane, causing accumulation of large clathrin-containing ‘scars’ on the cell surface. Intriguingly, the requirement for MYO7B in endocytosis is only restricted to α-Syn PFF and other cargos that enter cells via HSPGs. Thus, by identifying new regulatory factors for α-Syn PFF endocytosis, our study defines a mechanistically distinct clathrin-mediated endocytosis pathway that requires additional force generated by MYO7B and actin filaments.

**Significance:** The spreading of misfolded protein aggregates such as α-Synuclein preformed fibrils (α-Syn PFF) from cell to cell is a pathologic hallmark associated with the progression of many neurodegenerative diseases, but it is unclear how mammalian cells take up these large protein aggregates to initiate this prion-like protein transmission process. Here we define the mechanism of α-Syn PFF endocytosis using a combination of genetic, biochemical, and live-cell imaging techniques. Our study reveals how α-Syn PFF binds to the cell surface heparan sulfate proteoglycans using two lysine-bearing motifs and then enters cells following a Myosin-7B- and actin-dependent endocytosis mechanism that is specifically tailored for polycation-bearing cargos.

---

In Parkinson’s disease (PD), the major proteinaceous aggregate known as Lewy body is mostly comprised of a synaptic protein named α-Synuclein (α-Syn) (1). Coincidentally, genetic mutations in α-Syn-encoding gene were identified as a major contributing factor for Parkinson’s disease (2, 3). α-Syn is a small polypeptide that is aggregation-prone. Clinical studies showed that α-Syn-positive Lewy bodies can spread in patient brains during disease progression (4). Furthermore, following tissue transplantation therapy, grafted tissues can accumulate Lewy body-like aggregates over time, which does not occur if these cells are left intact in the donor (5, 6). These observations prompted the idea that α-Syn might undergo cell-to-cell transmission analogously to a prion protein (7, 8). Indeed, *in vitro* and animal studies showed that α-Syn preformed fibrils (α-Syn PFF) can be readily taken up by various cell types and subsequently transmitted to nearby cells that have not been exposed directly to α-Syn PFF (9–13).

At the cellular level, intercellular transmission of neurotoxic proteins like α-Syn comprises two biological events: release of α-Syn from donor cells via unconventional protein secretion and its internalization by recipient cells via endocytosis (14). The molecular mechanisms of these processes are poorly defined. Recent studies suggested nanotubes, exosome secretion, or a pathway termed misfolding-associated protein secretion (MAPS) as potential mechanisms for the release of monomeric and oligomerized α-Syn (15, 16). On the uptake side, α-Syn PFF can enter cells via clathrin-mediated endocytosis (CME) upon binding to a surface receptor(s) (17), but whether α-Syn endocytosis involves substrate specific endocytosis regulators has not been explored. This type of regulators may be particularly relevant to the development of PD drugs that target α-Syn endocytosis. Another unresolved issue is the identity of relevant membrane receptor(s) for α-Syn PFF, since several studies reported different candidates: heparan sulfate proteoglycans (HSPGs) and a membrane protein named LAG3 for unmodified α-Syn PFF (18, 19), and a glycoprotein named neurexin 1β for N-terminally acetylated α-Syn PFF (20).

HSPGs refer to a class of cell surface and extracellular matrix glycoproteins that carry one or more heparan sulfate (HS) chains bearing repeated disaccharide units. This modification was primarily found in the Syndecan and glycosylphosphatidylinositol (GPI)-anchored glypican families (21). The biosynthesis of HSPGs begins in the endoplasmic reticulum (ER), but its completion requires modification of the assembled sugar chains with sulfate using 3’-phosphoadenosine-5’-phosphosulfate (PAPS) as the sulfate donor, which occurs in the Golgi complex (22). Because HS chains carry negative charges, they can interact with a plethora of ligands on the cell surface, either mediating their uptake or activating downstream signal transduction (23, 24). The known HSPG ligands do not share any specific sequence motif for engaging HSPGs. Instead, the binding energy seems to be provided by electrostatic interactions, often involving positively charged residues from random sequences in ligands (21). In addition to α-Syn PFF, HSPGs have been implicated in the uptake of Tau PFF and Aβ (18, 25, 26), but how HSPGs interact with these neurotoxic protein aggregates is unclear.

In this study, we used an unbiased CRISPR/Cas9 screen to identify factors that regulate α-Syn PFF endocytosis in mammalian cells. The screen isolated known regulators (e.g., HSPGs), but also identified factors that had not been previously linked to α-Syn PFF uptake (an unconventional Myosin and actin). Mechanistically, our study defines the interaction of HSPGs with α-Syn PFF using biochemical assays and structural modeling. Importantly, we reveal a unique CME mechanism for α-Syn PFF and other polycation-bearing cargos, which unlike conventional CME, requires Myosin 7B (MYO7B) and actin filaments.

## Results

### A CRISPR screen identifies proteins involved in α-Syn PFF endocytosis

To dissect the mechanism of α-Syn PFF endocytosis, we labeled purified α-Syn with either a stable fluorophore (Alexa_594_) or a pH-sensitive dye (pHrodo). The pHrodo dye becomes activated only upon its arrival at the acidic late endosome/lysosome compartment, emitting red fluorescence (SI Fig. S1A). We used labeled proteins to prepare α-Syn PFF following a well-established protocol (27). Electron microscopy (EM) analysis of sonicated α-Syn PFF revealed particles of 20 to 100 nm in length (SI Fig. S1B), consistent with the reported α-Syn PFFs bearing pathogenic activities (27). We tested α-Syn PFF uptake in various cell types and found that both neuronal and non-neuronal cells could efficiently internalized α-Syn PFF (SI Fig. S1C), suggesting that the core endocytosis machinery for α-Syn PFF is not specific to neurons. Treating cells with dynamin inhibitors such as Dynasore (28) or Dynole 34-2 (29) significantly reduced α-Syn PFF uptake (SI Fig. S1D, E). Additionally, α-Syn PFF uptake was also dramatically attenuated in HeLa cells lacking the σ2 subunit of AP-2 (encoded by the *AP2S1* gene), a cargo adaptor in CME (SI Fig. S1F). These findings confirm CME-dependent α-Syn PFF endocytosis (17, 30).

To search for α-Syn PFF endocytosis regulators, we performed a pooled CRISPR/Cas9 screen (Figure 1A). We chose HEK293T cells because their fast-growing and high DNA recombination properties made them well suited for a CRISPR screen. Since pooled CRISPR screens often suffer from poor specificity and reproducibility, we took a strategy akin to the classical genetics approach. We treated HEK293T cells with a lentiviral GeCKO library expressing sgRNAs targeting all human genes. We then incubated mutagenized cells with pHrodo-labeled α-Syn PFF. Florescence-activated cell sorting (FACS) was used to sort α-Syn PFF negative cells into 96-well plates at single-cell resolution. When these cells formed colonies, we re-screened them to identify cells defective in α-Syn PFF uptake. From 1,248 clones screened, one (C10) showed almost no α-Syn PFF uptake (SI Fig. S2A), whereas 27 other clones showed reduced α-Syn PFF uptake. In this study, we characterized two clones, which revealed an unexpected requirement for the endocytosis of α-Syn PFF and other polycation-bearing cargos.

**Figure 1.**
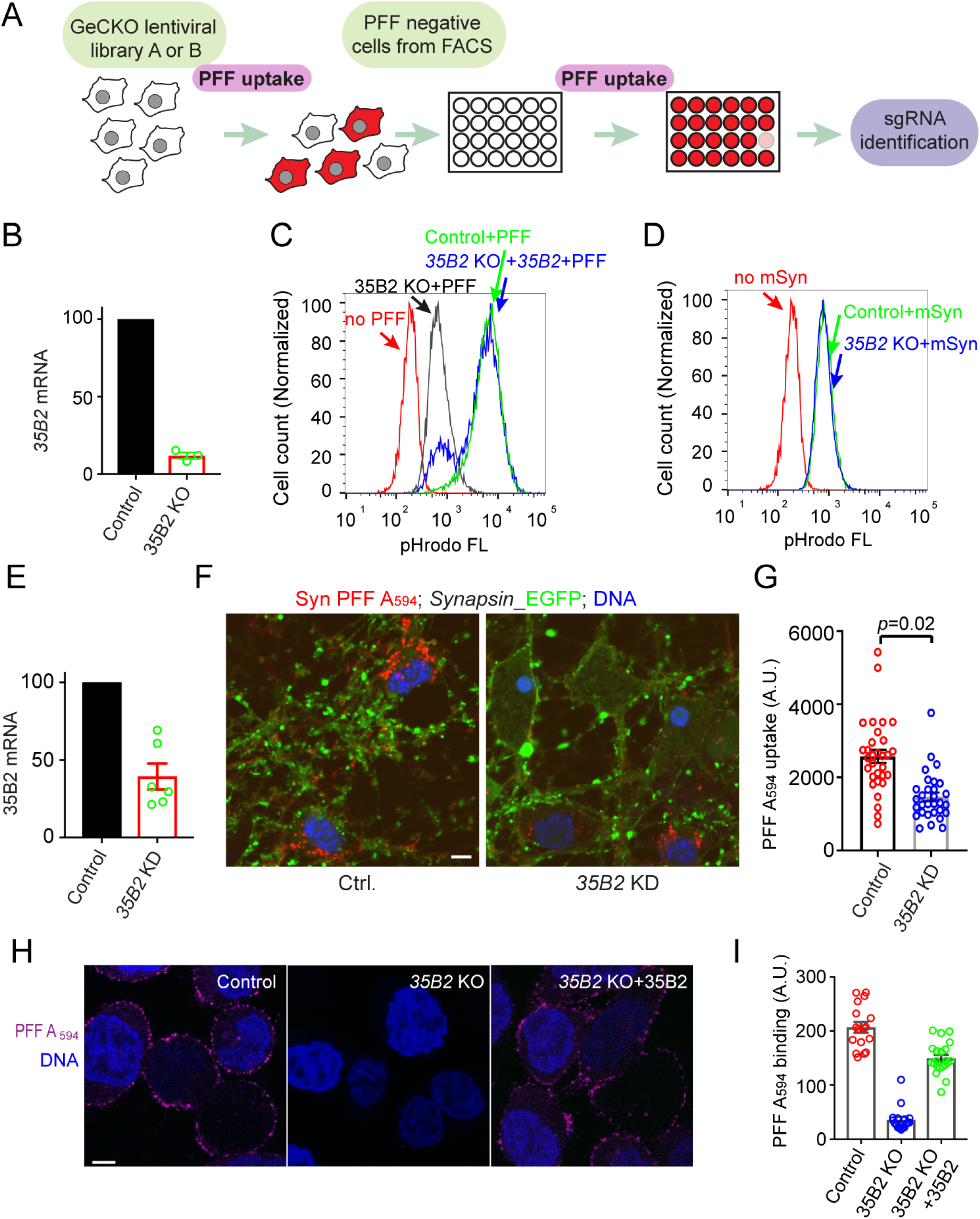
SLC35B2 is required for endocytosis of α-Syn PFF. **(A)** The CRISPR screen strategy. **(B-D)** *SLC35B2*-KO cells are defective in endocytosis of α-Syn PFF but not monomeric α-Syn. **(B)** Validation of *SLC35B2* (35B2)-KO cells by qRT-PCR. mRNA levels were normalized to that in control cells. Error bar, SEM, n=3. **(C)** Control, *SLC35B2*-KO, or *SLC35B2*-KO cells stably expressing FLAG-SLC35B2 were incubated with pHrodo-labeled α-Syn PFF (200nM for 4 h) and analyzed by FACS. FL, fluorescence. **(D)** Control and *SLC35B2*-KO cells were treated with labeled α-Syn monomer at 800nM overnight prior to FACS analysis. **(E-G)** Knockdown (KD) of SLC35B2 attenuates α-Syn PFF uptake in primary neurons. **(E)** Primary neurons were infected with lentivirus expressing the indicated shRNAs together with EGFP (driven by the Synapsin promoter) at days-in-*vitro 3* (DIV3). mRNA was purified from cells at DIV8 for qRT-PCR analysis. Error bar, SEM, n=2 biological repeats, each with triplicated PCR analyses. **(F)** Control or *SLC35B2*-KD neurons expressing EGFP (green) at DIV8 were incubated with α-Syn PFF Alexa_594_ (200nM) (red) for 4 h, stained with DAPI (blue), and analyzed by confocal microscopy. Scale bar, 5µm **(G)** Quantification of α-Syn PFF level in individual cells (indicated by dots) from two independent experiments. Error bar, SEM, *p* value from two-tailed t-test. A.U. arbitrary unit. **(H, I)** SLC35B2 is required for α-Syn PFF binding to the plasma membrane. (**H**) Control, *SLC35B2*-KO, or *SLC35B2*-KO cells re-expressing WT SLC35B2 were incubated with α-Syn PFF Alexa_594_ (200nM) (magenta) on ice for 30min, stained with DAPI (blue), and imaged by confocal microscopy. Scale bar, 5µm. The graph in **(I)** shows the quantification of α-Syn PFF surface level in individual cells from two independent experiments. A.U. arbitrary unit.

### HSPGs are required for α-Syn PFF endocytosis

Sequencing identified a sgRNA in one of the clones (C10), targeting *SLC35B2*. Analysis of *SLC35B2*-knockout (KO) cells confirmed a specific requirement for SLC35B2 in endocytosis of α-Syn PFF, but not monomeric α-Syn (mSyn) (Figure 1B-D). Furthermore, shRNA-mediated knockdown of SLC35B2 in primary neurons also reduced α-Syn PFF uptake (Figure 1E-G), but the uptake defect was less dramatic compared to that in *SLC35B2* KO cells, likely because of partial gene silencing. The *SLC*3*5B2* gene contains 4 exons. The identified sgRNA targets a sequence in exon 2 (SI Fig. S2B). SLC35B2 was known as a Golgi-localized membrane protein, required for transporting 3-phosphoadenosine 5’-phosphosulfate (PAPS) into the Golgi complex for protein sulfation (SI Fig. S2C) (31). We confirmed the Golgi localization of SLC35B2 by immunostaining with a FLAG antibody *SLC35B2*-KO cells stably expressing FLAG-tagged SLC35B2 (SI Fig. S2D). The functionality of ectopically expressed SLC35B2 was confirmed by its ability to rescue α-Syn PFF uptake in *SLC35B2*-KO cells (Figure 1C). Collectively, these results suggested a role for Golgi-localized protein sulfation in α-Syn PFF endocytosis.

Cell surface binding experiments showed that SLC35B2 was required for the recruitment of α-Syn PFF to the plasma membrane (Figure 1H, I). Given the known role of SLC35B2 in biosynthesis of HSPGs (31) and because recent studies suggested HSPGs as a potential receptor for α-Syn PFF (18, 32), we presumed that α-Syn PFF failed to bind *SLC35B2*-KO cells because these cells lacked HSPGs. Indeed, immunostaining with an antibody to HSPGs stained the plasma membrane of WT, but not that of *SLC35B2*-KO cells (SI Fig. S2E, F). Furthermore, KO of XYLT2, another key HSPG biosynthetic enzyme (33) diminished both cell surface HSPG signal (SI Fig. S2G) and α-Syn PFF binding (SI Fig. S2H, I). Thus, our unbiased genetic screen confirms HSPGs as critical mediators of α-Syn PFF-membrane interaction, which is essential for α-Syn PFF endocytosis.

### Electrostatic interactions recruit α-Syn PFF to the plasma membrane

To better characterize the α-Syn PFF-HSPG interaction, we developed a fluorescence probe that can track HSPGs in live cells. Since the oligosaccharide chains in HSPGs are enriched in negative charges (23), we reasoned that an engineered GFP variant carrying net 36 net positive charges (GFP+) (34) might be a specific probe for HSPGs. Indeed, several lines of evidence support this notion. First, live-cell confocal microscopy showed that GFP+, but not a GFP variant bearing extra negative charges (GFP-) bound efficiently to the cell surface, initially forming a smooth profile, which was rapidly converted to a punctate pattern reminiscent of cells treated with α-Syn PFF Alexa_594_ (Figure 2A). The interaction of GFP+ with the cell surface was completely dependent on SLC35B2 as no GFP+ staining was detected in *SLC35B2*-KO cells (Figure 2B, panel 2 vs. 1, 3). Furthermore, when cells were pre-treated with GFP+, cell surface HSPGs could not be stained by antibodies to HSPGs (Figure 2C), suggesting that GFP+ competes with HSPG antibodies for the same binding site. Because cells treated with GFP+ and α-Syn PFF Alexa_594_ showed extensive co-localization of these two cargos on the cell surface (Figure 2D), and because pre-treating α-Syn PFF with GFP-reduced the binding of α-Syn PFF to cells (Figure 2E), electrostatic interactions must provide the major energy that recruits α-Syn PFF to the cell surface (Figure 2F).

**Figure 2.**
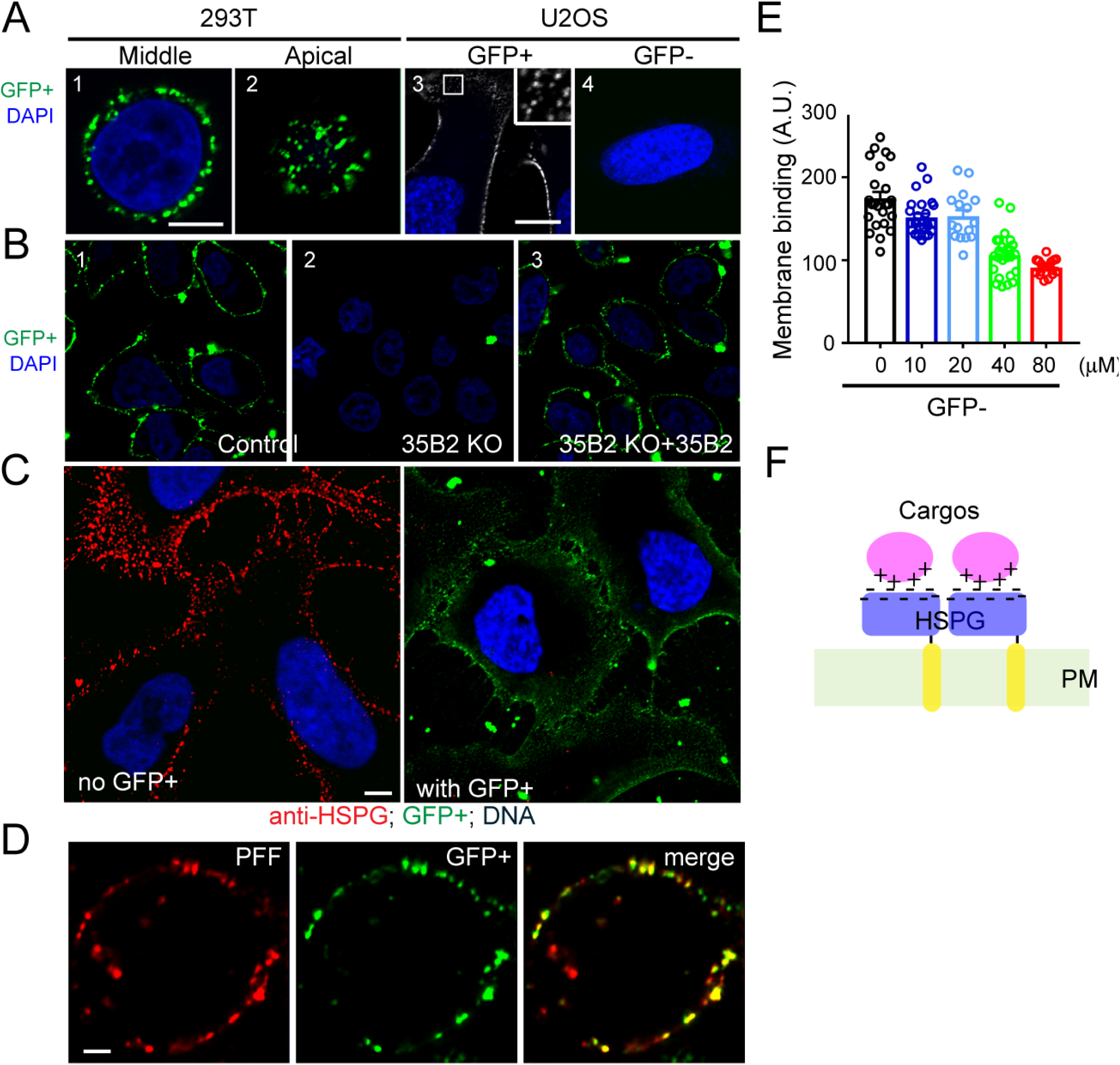
α-Syn PFF interacts with cells via electrostatic interactions. **(A)** GFP+ binds to the plasma membrane. Panels 1-2 show two confocal sections of a HEK293T cell stained with 200nM GFP+ (green) and DAPI (blue) for 5min. Scale bar, 5µm. Panels 3-4 show U2OS cells incubated with 200nM of GFP+ (Panel 3) or GFP- (Panel 4) and then stained with DAPI. The inset shows an enlarged view of the box in panel 3. Scale bars, 10µm. **(B)** The interaction of GFP+ with the plasma membrane depends on SLC35B2. Control (Panel 1) or *SLC35B2*-KO HEK293T cells with (Panel 3) or without (Panel 2) re-expression of WT SLC35B2 were stained with GFP+ and DAPI. Scale bar, 5µm. **(C)** GFP+ inhibits the binding of HSPG antibody to the cell surface. U2OS cells were either pre-treated with GFP+ (200nM) (Right) for 10min or left untreated (Left) and then stained with antibodies to HSPGs (red) and DAPI (blue). Scale bar, 5µm. **(D)** Co-localization of GFP+ with α-Syn PFF in cells. Cells incubated with α-Syn PFF Alexa_594_ together and GFP+ were imaged. **(E)** GFP-attenuates α-Syn PFF binding to the plasma membrane. Cells were incubated with α-Syn PFF Alexa_594_ (200nM) in the presence of the indicated concentrations of GFP-. α-Syn PFF binding to the cell surface was quantified by confocal imaging. **(F)** An HSPG-cargo interaction model.

### Two K-T-K motifs in oligomerized α-Syn enable HSPG interactions

Previous studies showed that two α-Syn monomers could form either a face-to-face or back-to-back dimer. Many dimers then stack together to form long fibrils of “rod” or “twisted” conformers (35, 36). To elucidate how these α-Syn PFF conformers interact with HSPGs, we used the ClusPro program to model the interaction of α-Syn PFF conformers with heparin, a heparan sulfate analog that competitively inhibits the binding of α-Syn PFF to cells (18). The modeling resulted in 14 clusters, falling into 3 groups as shown in Figure 3A and SI Figs. S3A, B, respectively. Interestingly, in all clusters, the interactions between α-Syn PFF and heparin were mediated by two K-T-K motifs (K43-T-K45 and K58-T-K60) in α-Syn fibrils. In the twisted conformer, each of these lysine-rich motifs lined up to form a shallow concave surface that accommodate a di-heparin molecule bearing six sulfate groups (Figures S3A, B). In contrast, in the rod conformation, two K-T-K motifs from distinct α-Syn molecules in a dimeric fibril joined together to form one deep binding groove for di-heparin (Figure 3A, B). As a result, each fibril could only offer two binding grooves, but within each binding site, the heparin molecule was sandwiched by four rows of lysine residues, forming extensive electrostatic interactions as well as hydrogen bonds with α-Syn (Figure 3C, SI Fig. S3C). Thus, α-Syn fibrils in the rod conformer is expected to have a much higher affinity for HSPGs than the twisted conformer. These models explain why HSPGs is only required for the uptake of α-Syn fibrils but not monomeric α-Syn. The result also suggests that a twisted α-Syn oligomer containing three protomers is the minimum requirement for binding one heparin unit; in contrast, for the rod conformer, five to six α-Syn protomers are required to form a higher affinity heparin binding site.

**Figure 3.**
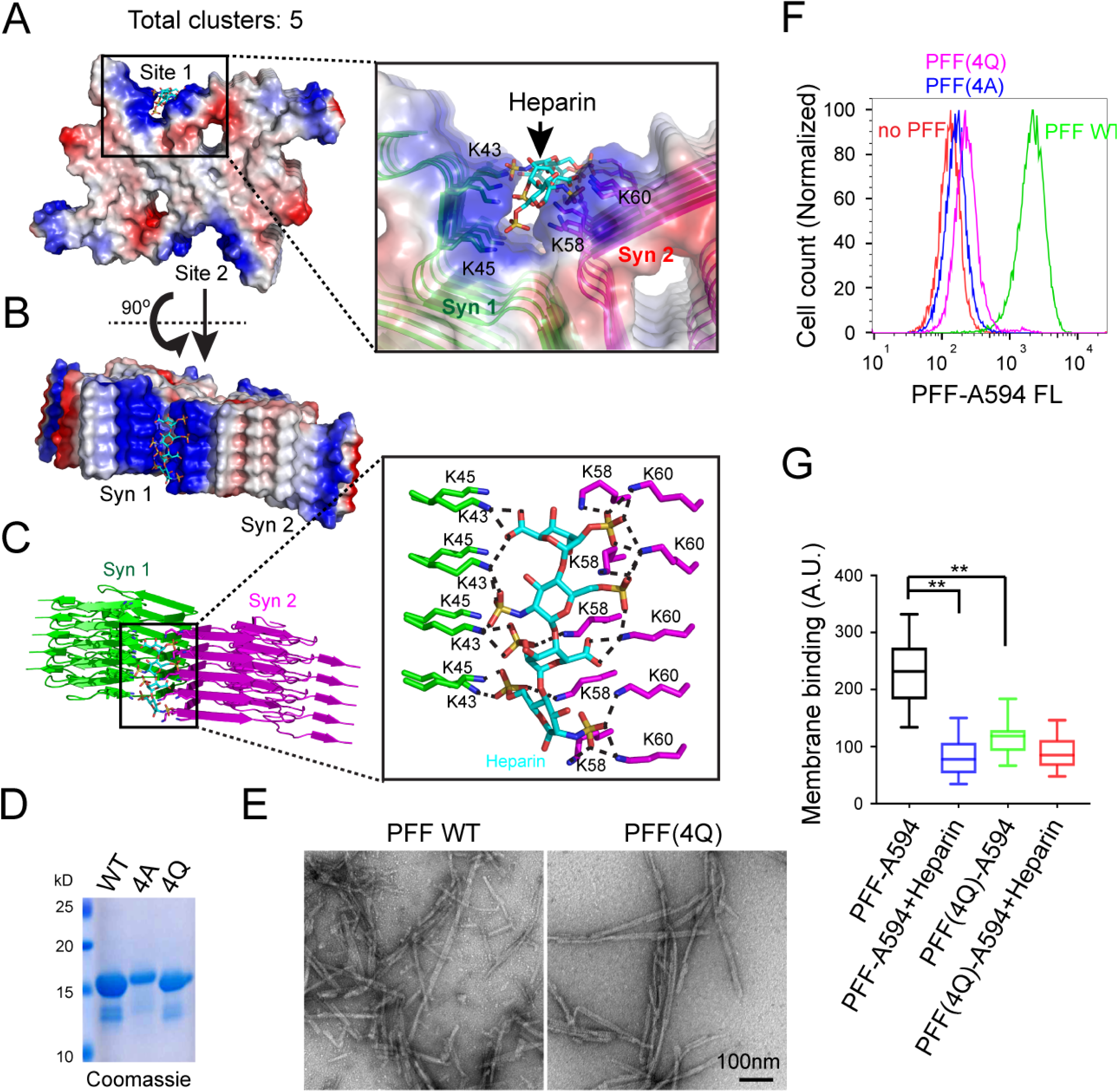
Two K-T-K motifs in α-Syn PFF enable specific interactions with heparin. **(A-C)** A model of the α-Syn PFF rod conformer in complex with heparin. **(A, B)** Shown is a model representing 5 clusters. In this model, each α-Syn PFF dimeric fibril contains two heparin binding pockets (Site 1 and Site 2) that each form a long grove to accommodate a heparin molecule. **(C)** Heparin is sandwiched by four rows of lysine residues in α-Syn fibril. Dashed lines in **(C)** label complementarily charged atoms within 3 Å. **(D)** Purified WT α-Syn and charge mutants (4A and 4Q) analyzed by SDS-PAGE and Coomassie blue staining. **(E)** Representative negative stained EM images show PFFs formed by WT α-Syn and the 4Q α-Syn mutant. **(F, G)** α-Syn PFF Alexa_594_ formed by the charge mutants are defective in endocytosis **(F)** and cell surface binding **(G)**. **(F)** HEK293T cells were treated with the indicated α-Syn PFF Alexa_594_ for 3 h before FACS analysis. The whisker plot in **(G)** shows the relative level of α-Syn PFF binding to cells as determined by confocal imaging of U2OS cells treated with the indicated α-Syn PFF Alexa_594_ at 400nM in the presence or absence of heparin (4µM) on ice. **, p<0.01, by two-tailed Student’s t-test.

To validate our models, we generated α-Syn mutants with the four lysine residues (K43, K45, K58, and K60) substituted to either glutamine (4Q) or alanine (4A) (Figure 3D). These mutants could form long fibrils similarly to WT α-Syn (Figure 3E). However, compared to WT α-Syn PFF, the uptake of these mutant α-Syn PFFs was dramatically reduced (Figure 3F). By contrast, the uptake of monomeric α-Syn was not affected by these mutations (SI Fig. S3D). Furthermore, cell binding experiments showed that the interaction of α-Syn PFF with the plasma membrane was significantly inhibited by these mutations (Figure 3G). These results strongly suggest that α-Syn binds HSPGs via a specific sequence motif that forms binding sites only upon α-Syn oligomerization.

### HSPG-mediated endocytosis requires MYO7B and actin filaments

Using a similar approach, we identified a *MYO7B*-specific sgRNA from a clone partially defective in α-Syn PFF endocytosis. Like for *SLC35B2*, the sgRNA targeting *MYO7B* mapped to an early exon (Figure 4A). Reconstructed *MYO7B*-KO HEK293T cells or cells treated with *MYO7B*-specific siRNA showed that depletion of MYO7B diminished endocytosis of α-Syn PFF (Figure 4B, C, SI Fig. S4A), while re-expression of EGFP-MYO7B in *MYO7B*-KO cells largely rescued the α-Syn PFF uptake phenotype (SI Fig. S4B). An α-Syn PFF uptake defect was similarly observed in primary neurons expressing *MYO7B*-specific shRNA (Figure 4F-H). Strikingly, the function of MYO7B in endocytosis appeared to be restricted to HSPG-dependent cargos because like HSPG-deficient cells (23), *MYO7B*-KO cells were also defective in uptake of GFP+ (SI Fig. S4C, D) and of DNA in complex with a polycation carrier (SI Fig. S4E). Furthermore, HSPG-dependent uptake of a lentiviral reporter expressing GFP was significantly reduced in *MYO7B-*KO cells (SI Fig. S4F, G). On the other hand, endocytosis of monomeric α-Syn or transferrin was not affected in *MYO7B*-KO cells (Figure 4D, E).

**Figure 4.**
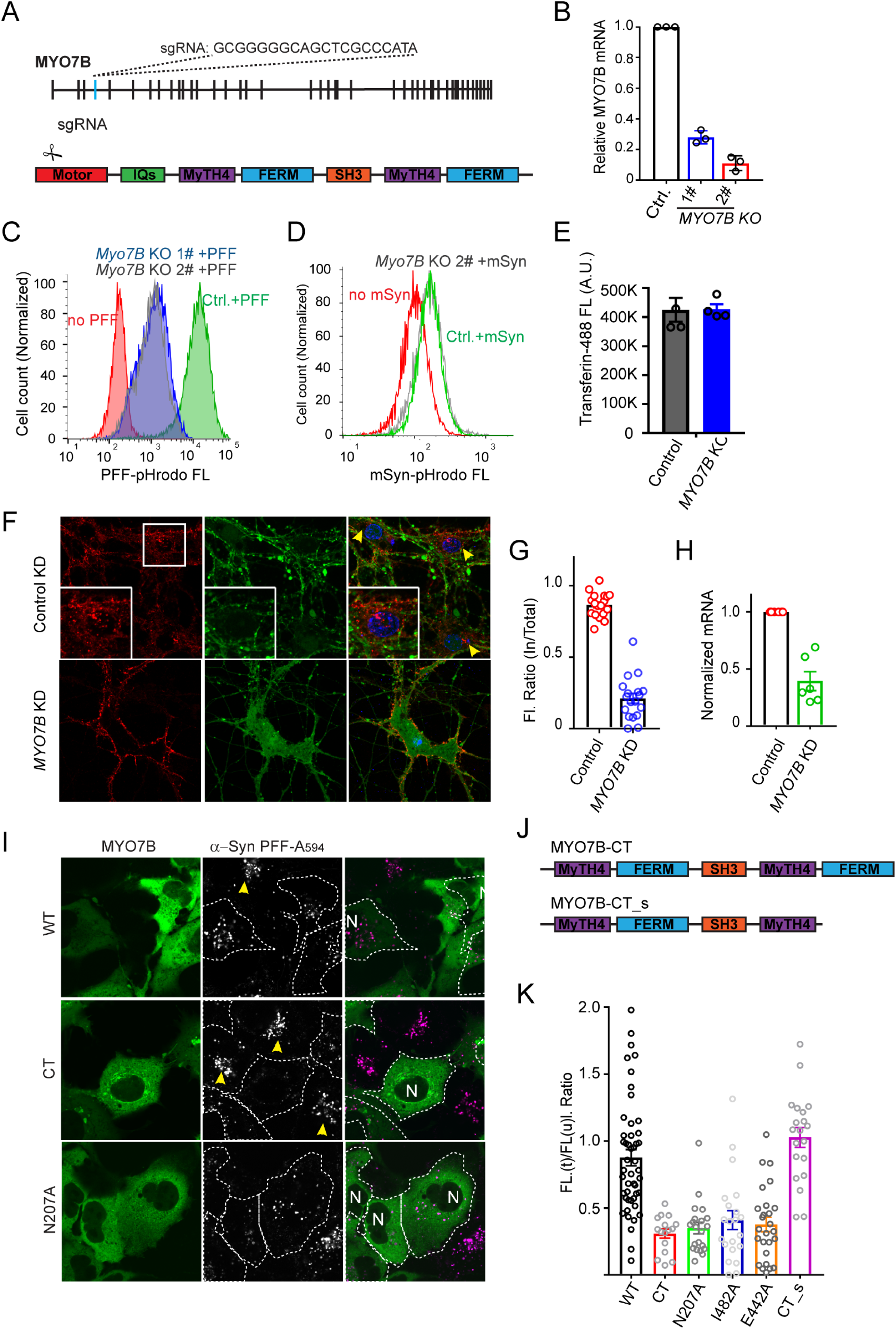
HSPG-mediated endocytosis requires MYO7B and actin filaments. **(A)** Mapping the identified MYO7B sgRNA. **(B)** Validation of *MYO7B*-KO cells by qRT-PCR. mRNAs extracted from two clones of *MYO7B* KO cells and a control (Ctrl.) clone were analyzed by qRT-PCR. Error bars, SEM, n=3. **(C)** *MYO7B* KO cells are defective in α-Syn PFF uptake. Control (Ctrl.) and *MYO7B*-KO clones were incubated with pHrodo-labeled α-Syn PFF for 4 h before FACS analyses. **(D)** MYO7B is not required for endocytosis of α-Syn monomer. Control or *MYO7B*-KO cells were incubated with α-Syn monomer (800nM) overnight prior to FACS analysis. **(E)** MYO7B is not required for transferrin uptake. Control or *MYO7B*-KO cells were incubated with fluorescein-labeled transferrin (50µg/ml, 3h), washed, and analyzed with a fluorometer. Error bars, SEM, n=3. **(F-H)** KD of *MYO7B* in primary neurons reduces α-Syn PFF endocytosis without affecting its binding to the plasma membrane. **(F)** Primary neurons infected with the indicated shRNA-expressing lentivirus together with *Synapsin_EGFP* lentivirus at days-in-*vitro 3* (DIV3) were incubated with α-Syn PFF Alexa_594_ (200nM) for 4 h and imaged at DIV8. The insets show an enlarged view of the box. Arrows indicate peri-nuclear enrichment of α-Syn PFF positive vesicles in control cells. The graph in **(G)** shows the ratio of internalized α-Syn PFF relative to total α-Syn PFF in individual cells. **(H)** qRT-PCR evaluation of MYO7B mRNA levels using the same batch of cells. Error bars, SEM, n=two biological repeats with triplicated PCR analyses. **(I-K)** MYO7B dominant negative mutants inhibit α-Syn PFF uptake. **(I)** Representative confocal images of COS7 cells transfected with the indicated MYO7B plasmids and treated with α-Syn PFF Alexa_594_ (400nM) for 3 h. Arrows indicate normal α-Syn PFF uptake in untransfected cells, which was used to normalize uptake. N, nuclei. **(J)** The graph shows the domain structure of the truncated MYO7B mutants. **(K)** Quantification of α-Syn PFF fluorescence (FL) in individual cells transfected (t) with the indicated MYO7B mutants normalized by the signal in untransfected cells (u).

To test whether HSPG-mediated endocytosis involves the motor activity of MYO7B, we ectopically expressed MYO7B mutants that either lack the motor domain (CT) or carry a mutation that disrupts the ATPase cycle of the motor domain (N207A, I482A, E442A) (37). If coupling the motor activity to cargo binding was required for α-Syn PFF uptake, these mutants should inhibit α-Syn PFF uptake in a dominant negative manner. Indeed, cells expressing these mutants showed significantly reduced α-Syn PFF uptake compared to neighboring untransfected cells or cells transfected with WT MYO7B (Figure 4I-K). Furthermore, deleting the C-terminal membrane-binding FERM domain abolished the dominant negative activity of MYO7B-CT, suggesting that membrane binding by a motor-defective MYO7B mutant accounts for the observed endocytosis defect. These results, together with the observation that actin-binding compounds (e.g. Jasplakinolide and Latrunculin A) that interfere with the function of F-actin also inhibited α-Syn PFF endocytosis (SI Fig. S5A, B), strongly suggest that optimal α-Syn PFF endocytosis requires a membrane-associated functional interplay between MYO7B and actin filaments.

### MYO7B maintains membrane dynamics at clathrin-enriched membrane domains

To elucidate the mechanism of MYO7B-mediated endocytosis, we used total internal reflection fluorescence microscopy (TIRF) to examine the localization of MYO7B in U2OS cells stably expressing mCherry-tagged clathrin light chain (mCh-CLC) to label clathrin-coated pits (CCPs) together with either GFP-WT MYO7B or GFP-MYO7B-CT. Consistent with the reported localization of MYO7B to plasma membrane-derived microvilli (37), GFP-MYO7B also bound to the plasma membrane in U2OS cells, forming discrete dots (Figure 5A). Time-lapse microscopy showed that MYO7B-positive signals fluctuated rapidly, displaying a flickering pattern (Figure 5B, SI movie1). Interestingly, the MYO7B-CT mutant appeared to form more stable interactions with the membranes (Figure 5A), resulting in bright foci that remained constant for a long period of time (SI movie2). We concluded that the MYO7B C-tail domain can bind to the plasma membrane, while its actin-binding domain contributes to the dynamic property of this interaction.

**Figure 5.**
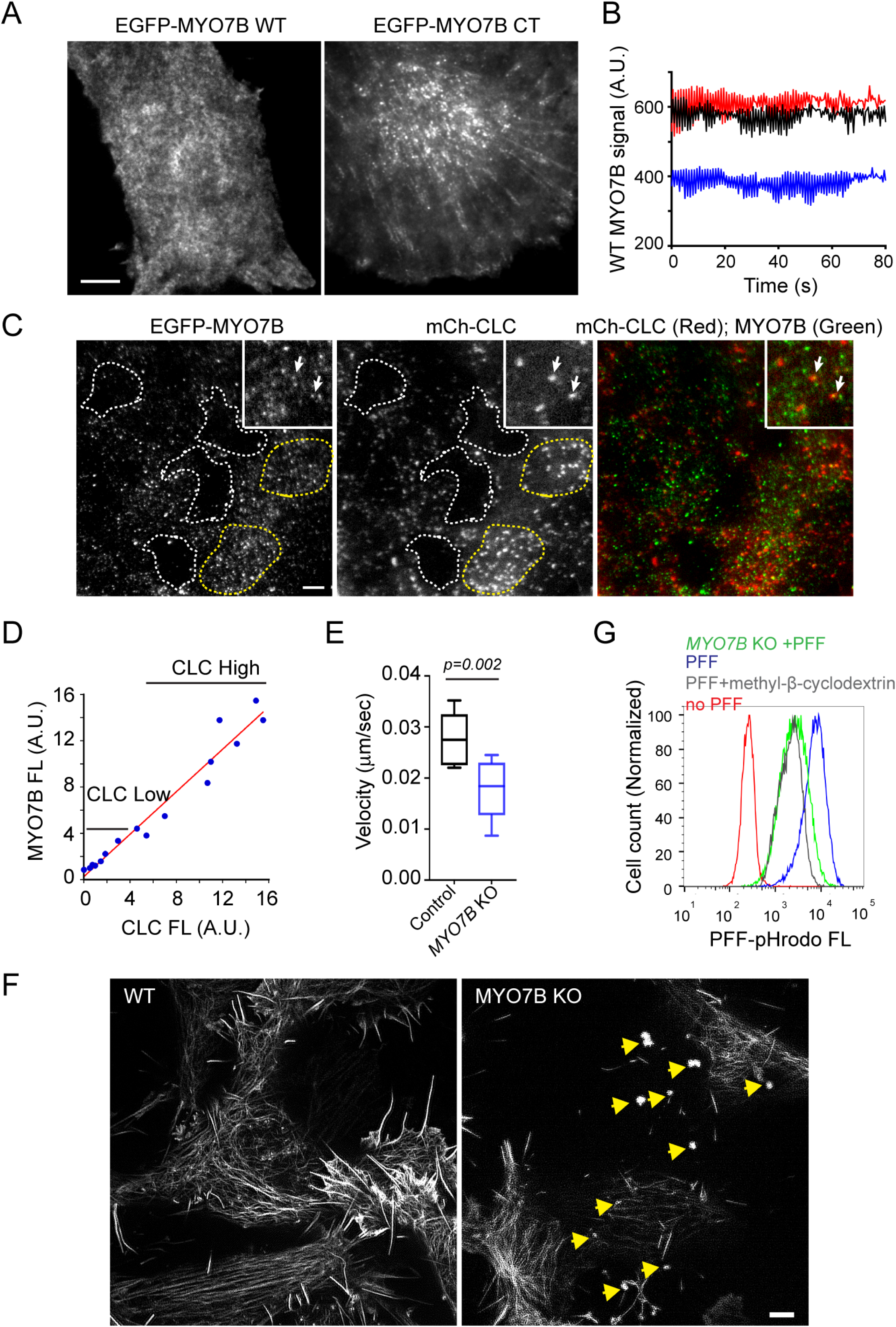
MYO7B maintains plasma membrane dynamics at domains enriched in clathrin. **(A)** TIRF microscopy analysis of plasma membrane binding by EGFP-WT MYO7B and EGFP-CT MYO7B in cells stably expressing low level of these proteins. Scale bar, 5µm. **(B)** Quantification of representative EGFP-WT MYO7B signal on the plasma membrane in a time-lapse video. Shown are fluorescence intensity of three randomly chosen spots measured by Fiji. A.U., arbitrary unit. **(C, D)** MYO7B binds the plasma membrane at clathrin-enriched domains. **(C)** TIRF microscopy analysis of cells stably expressing EGFP-WT MYO7B and mCherry-tagged clathrin light chain (mCh-CLC). The insets show a close-up view of a cell, revealing limited colocalization between MYO7B and CLC. The dashed lines indicate surface regions either lacking clathrin (white) or enriched in clathrin (yellow). The graph in **(D)** shows the correlation between CLC and MYO7B fluorescence (FL) intensity in cell surface domains as outlined in Figure 5C (R^2^=0.956 by linear regression). **(E)** *MYO7B* inactivation affects plasma membrane dynamics. Control or MYO7B-KO cells were stained with GFP+ and imaged by time-lapse confocal microscopy (SI Fig. S5C and SI movie3 and 4). The whisker graph shows the plasma membrane ruffling velocity determined by Nikon Element analysis of videos taken from three independent experiments. Control, n=6 cells, MYO7B-KO, n=12 cells. *p* value by two-tailed unpaired Student’s t-test. **(F)** *MYO7B*-KO cells have reduced actin fibers under the plasma membrane. TIRF-SIM analysis of WT or *MYO7B-*KO cells transfected with EGFP-Tractin. Note the presence of actin-containing aggregates in *MYO7B*-KO cells (indicated by arrows). Scale bar, 10µm. **(G)** Depletion of cholesterol inhibits α-Syn PFF endocytosis. Cells were treated with methyl-β-cyclodextrin (5mM) or DMSO for 30min prior to incubation with α-Syn PFF pHrodo and FACS analysis.

When co-localization of MYO7B with CCPs was examined by two-color TIRF, we found limited overlapping signals (Figure 5C, insets). However, we noticed that MYO7B was not uniformly distributed on the plasma membrane. Instead, certain areas had more MYO7B dots than others (Figure 5C). Intriguingly, MYO7B-high plasma membrane domains were often enriched in CCPs (Figure 5C, D). We therefore postulated that dynamic interactions of MYO7B within certain plasma membrane domains enable membrane flexibility and deformability, which might in turn facilitate the maturation of CCPs located in the same membrane domain.

To measure plasma membrane dynamics, we used GFP+ to label the cell surface and then took time-lapse videos. In WT cells, the plasma membrane usually undergoes dynamic expansion and retraction, generating a ruffling motion (SI Fig. S5C; SI movie3). By contrast, the membranes of *MYO7B*-KO cells had significantly reduced mobility (SI movie4, Figure 5E). Furthermore, TIRF microscopy showed that *MYO7B-*KO cells had a significantly reduced amount of actin filaments underneath the plasma membranes (Figure 5F). These results suggest that MYO7B may regulate actin assembly to maintain membrane dynamics.

If MYO7B promotes endocytosis via membrane dynamics at cargo binding sites, other agents capable of increasing membrane stiffness should similarly inhibit α-Syn PFF uptake. To test this idea, we treated cells with the cholesterol-depleting drug methyl-β-cyclodextrin, which increases membrane stiffness in mammalian cells (38). Consistent with our hypothesis, methyl-β-cyclodextrin also reduced α-Syn PFF uptake similarly as depletion of *MYO7B* (Figure 5G).

### MYO7B facilitates the maturation of clathrin-coated pits

Since α-Syn PFF endocytosis requires CME, we characterized the effect of *MYO7B* inactivation on CCP morphology using TIRF microscopy. In WT cells transfected with GFP-CLC, TIRF detected small CLC-positive fluorescence puncta at the basal membranes (Figure 6A, panel 1). In time-lapse videos, these CLC-positive puncta frequently detached from the membranes, vanishing into the cytosol due to endocytosis (Figure 6B, upper panels). Intriguingly, in *MYO7B*-KO cells, only a fraction of CCPs behaved like those in WT cells, while in ∼80% of the cells, we detected large CCP clusters that were completely immobile (Figure 6A, panel 2; 6B, lower panels). The CCP clustering phenotype was also observed in GFP-CLC stable U2OS cells transfected with a *MYO7B*-specific siRNA (Figure 6A, panel 4 vs. 3). Additionally, cells treated with Jasplakinolide, Dynole-34-2, or methyl-β-cyclodextrin (MBCD) also accumulated clustered CCP puncta at the basal membranes, coincident with α-Syn PFF endocytosis inhibition (Figure 6C, D). These results suggest that MYO7B-dependent membrane dynamics might faciliate CCP function in HSPG-mediated endocytosis. When CCPs fail to mature, these endocytosis-defective pits accumulate at the basal plasma membranes and form clusters.

**Figure 6.**
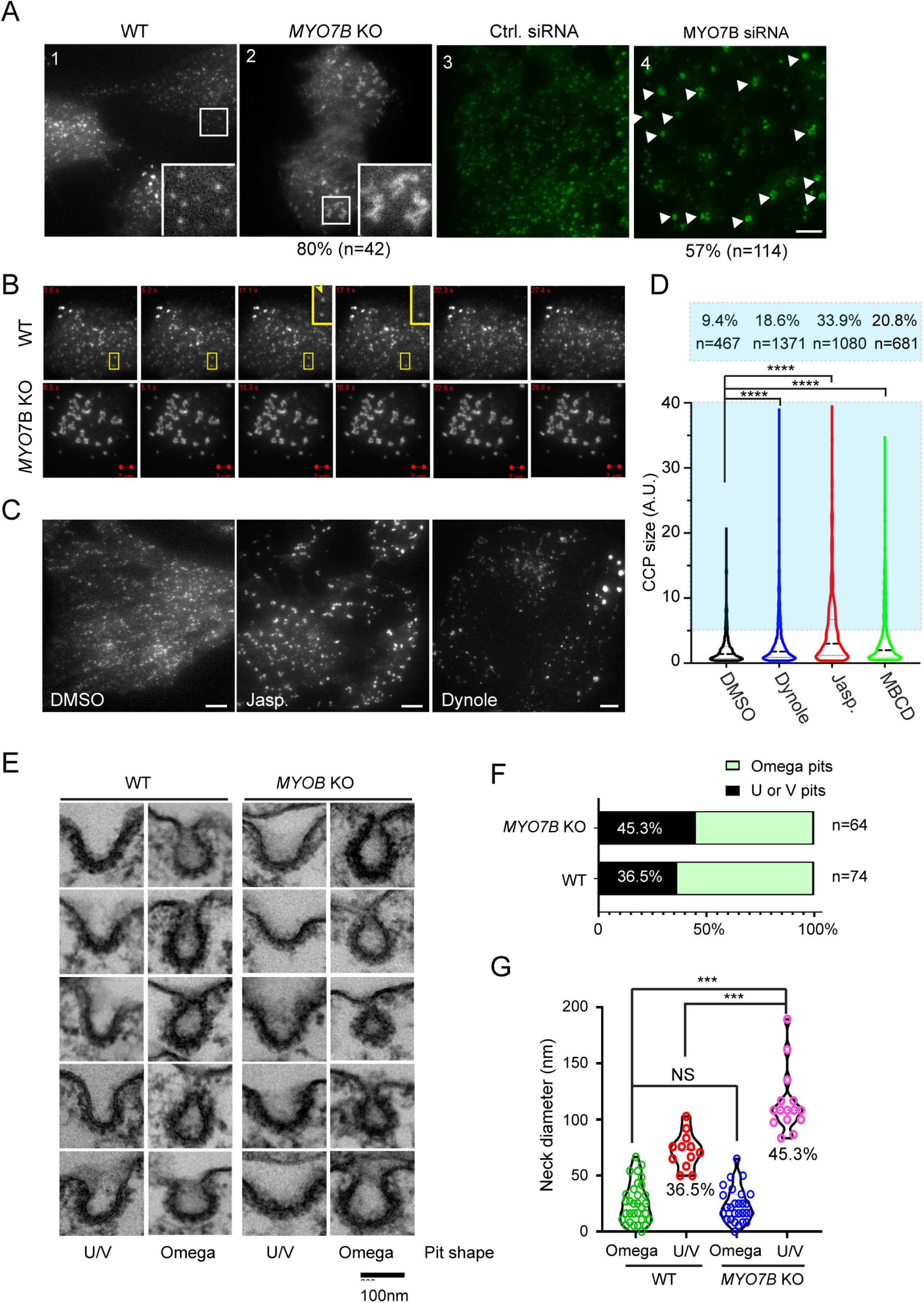
MYO7B inactivation affects the maturation of CCPs. **(A, B)** MYO7B depletion causes CCPs to cluster. **(A)** TIRF microscopy analysis of CCPs in Control (panel 1) or *MYO7B*-KO (panel 2) HEK293T cells transiently transfected with GFP-CLC, or in GFP-CLC stable U2OS cells transfected with control (panel 3) or MYO7B (panel 4) siRNA. ∼80% of *MYO7B*-KO cells show the phenotype shown in panel 2. Insets show enlarged views of the boxed regions. Scale bar, 5µm. **(B)** Images from time-lapse videos showing CCP dynamics on the cell surface. Boxes and insets show an example of CCP (arrow) vanishing into the cytosol in a WT cell. Scale bars, 2 µm. **(C, D)** The effect of Jasplakinolide (JASP), Dynole 34-2 (Dynole), and methyl-β-cyclodextrin (MBCD) on CCP morphology was analyzed by TIRF in GFP-CLC stable U2OS cells. **(C)** shows representative images of DMSO-, JASP- (100nM 1h), or Dynole- (18µM 2h) treated cells. Scale bar, 5 µm. The violin plot in **(D)** shows the relative CCP sizes under different drug-treated conditions. The numbers indicate the percent of clustered CCPs. N=number of pits analyzed, ****, *p* <0.0001, by one-way ANOVA. **(E-G)** Transmission EM analysis of CCP morphology in control and *MYO7B*-KO cells. Control or *MYO7B*-KO cells were incubated with α-Syn PFF (400nM, 1h), and then fixed for EM analysis. Images in **(E)** show representative CCPs. The graph in **(F)** shows the relative abundance of U- and V-shaped pits vs. Omega-shaped pits in control and *MYO7B*-KO cells. n=pit number. The violin plot in **(G)** shows the neck length of CCPs measured from two independent experiments. ***, p<0.001, by unpaired Student’s t-test. NS, not significant.

To test abovementioned hypothesis, we used transmission EM to examine the morphology of CCPs at the apical and lateral membranes where HSPG-mediated endocytosis occurs. To capture defects associated with α-Syn PFF uptake, we first incubated cells with α-Syn PFF at 37 °C for 1 h to initiate PFF uptake. EM analyses identified ∼4 CCPs per cell section in WT cells, but only 1 CPP per section on the plasma membrane of *SLC35B2*-KO cells (SI Fig. S6A, B), suggesting that the detected CCPs were mostly linked to HSPG-mediated endocytosis. Although the number of CCPs in *MYO7B*-KO cells were comparable to that of WT cells, we found that compared to WT cells, *MYO7B*-KO cells had ∼9% increase in the number of U- or V-shaped pits, and a concomitant reduction of Omega-shaped pits (Figure 6E, F). Additionally, for the U- or V-shaped pits, the average neck diameter was significantly increased in *MYO7B*-KO cells, whereas the neck diameter for Omega-shaped pits was comparable between WT and *MYO7B*-KO cells (Figure 6G). Since U- and V-shaped CCPs are thought to be precursors of Omega-shaped CCPs, these results support the notion that MYO7B is required for optimal CCP function.

## Discussion

### The role of HSPGs in endocytosis

HSPGs have been implicated in endocytosis of a variety of cargos (23), and the substrate diversity was attributed to non-specific electrostatic interactions between cargos and HSPGs. Consistent with this notion, in this study, we have shown that a GFP variant bearing net positive charges (GFP+) can be used as a probe to label HSPGs in mammalian cells. However, our study also shows that certain cargos like α-Syn PFF can interact with HSPGs via specific sequence motifs. Interestingly, among the 15 lysine residues in α-Syn, only 4 are specifically involved in interaction with HSPGs. These residues by themselves do not form the binding site, but when α-Syn assembles into stacked oligomers, they can line up to form long binding grooves that accommodate the linear oligosaccharide chains of HSPGs. Interaction with α-Syn PFF does not require the protein core of HSPGs, meaning that any of the 16 members of the HSPG family could possibly serve as a recruiter for α-Syn PFF. Importantly, our findings also suggest that the two previously established α-Syn PFF conformers have different affinities to HSPGs, which may correspond to the reported α-Syn PFF strains bearing different pathogenic activities (39, 40). Although we cannot exclude the involvement of other α-Syn motifs in cell surface binding, the identification of the K-T-K motifs in α-Syn as a major binding element for HSPGs may help design specific inhibitors that target α-Syn PFF entry for PD.

Despite its essential function in recruiting α-Syn PFF, HSPGs may not function as the sole receptor for α-Syn PFF because some HSPG family members do not have a cytoplasmic domain that is required for interacting with endocytosis effectors such as clathrin-coat components. Along this line, a recent study suggested LAG3 as another membrane receptor for α-Syn PFF in neuronal cells (19). We propose that HSPGs may cooperate with distinct membrane receptors in different tissues, which jointly mediate α-Syn PFF endocytosis. Recruitment of α-Syn PFF to the plasma membrane by HSPGs should increase its local concentration, and thus its avidity for a downstream receptor.

### A role for MYO7B and actin filaments in HSPG-mediated endocytosis

It is well established that mechanical force generated by actin polymerization can drive membrane invagination in CME in yeast and in mammalian cells bearing high membrane tension (41–43). However, the function of myosin motors in CME is much less clear, particularly in higher eukaryotes. In *S. cerevisiae*, the type I myosins (Myo3p and Myo5p) can promote the recruitment of actin assembly factors to the site of endocytosis, causing spatially-restricted actin polymerization to deform the membranes (44, 45). The mammalian class I Myosin-1E was also found at CCPs, coincident with a burst of actin assembly. However, depletion of Myosin-1E had no discernible effect in transferrin uptake under normal growth conditions (46). Another study suggested a role for a class VI myosin in CME based on its localization to CCPs/vesicles (47). However, a more recent study showed that myosin-6 is only localized to uncoated vesicles (48). On a related note, genetic studies have implicated MYO7A as a regulator of Listeria phagocytosis (49, 50), but this process is independent of CME. Thus, whether mammalian CME involves myosin(s) and if so, in what capacity has been unclear.

Our study now provides compelling evidence that MYO7B serves a specific function in CME, at least for certain polycation-bearing cargos that enter cells via HSPGs. Live-cell imaging showed that upon binding to the cell surface, GFP+ is rapidly clustered into patches (Figure 2), suggesting that HSPGs may undergo oligomerization upon ligand binding. Because membrane invagination during CME is expected to bring cargos and HSPGs close to each other, which would increase the repellent force between molecules bearing the same charge, we reason that MYO7B and actin filaments may provide the extra energy to overcome this membrane-bending barrier. Because actin was found both under the plasma membrane and on the surface of some α-Syn PFF-bearing endocytic vesicles (SI Fig. S6C), we propose that MYO7B either promotes actin assembly or stabilize actin filaments at membranes near oligomerized HSPG patches, which in turn facilitates membrane protrusions. These protrusions eventually wrap around cargos, forming Omega-shaped CCPs with actin filaments surrounding the CCP neck. Membrane severance by dynamin subsequently releases the vesicles into the cytoplasm, leaving an F-actin ‘scar’ attached to it (SI Fig. S6D). Additional studies will be required to reveal the mechanistic details of MYO7B-mediated CME as well as additional mechanisms that contribute to α-Syn PFF endocytosis. Nevertheless, our study has for the first time linked the function of MYO7B and actin filaments to HSPG-mediated endocytosis of α-Syn PFF and other polycation-bearing cargos. This unique feature may be exploited to develop drugs that prevent the transmission of α-Syn pathology in PD.

## Materials & Methods

### Cell lines and DNA transfection

All reagents are listed in SI table 1. HEK293T, U2OS, COS7, and HeLa cells were purchased from ATCC. HEK293FT cells were from Invitrogen. Cells were maintained in Dulbecco’s Modified Eagle’s Medium (DMEM, Corning) containing 10% fetal bovine serum (FBS) and antibiotics (penicillin/streptomycin, 10 U/ mL). All cell lines were maintained at 37 °C in a 5% CO_2_ humidified atmosphere. Cells stably expressing GFP- or mCherry-tagged clathrin light chain (CLC) or GFP-tagged MYO7B WT and the CT mutant were generated using U2OS cells. Specifically, these cells were seeded in a 6-well plate and transfected with the reporter constructs using Lipofectamine 2000. Two weeks post transfection, cells stably expressing the reporters were sorted out by FACS. Stable KO cells were generated by infecting HEK293T cells with lentivirus expressing sgRNAs targeting the gene of interest. Cells were selected by 0.35 µg/ml puromycin for 4 days to generate cells stably expressing the sgRNAs. Single cell-derived KO clones were obtained by sequential dilution or by FACS sorting.

Cell transfection was performed using TransIT-293 reagent (Mirus) for HEK293T, or Lipofectamine 2000 (Invitrogen) for COS7, HeLa and U2OS cells following the manufacturer’s instructions. For RNAi-mediated gene silencing, Lipofectamine RNAiMAX (Invitrogen) was used according to the manufacturer’s protocol. U2OS cells were seeded in 4 well chamber and transfected with 60 picomoles siRNA per well in antibiotic free medium for 48h before imaging analyses.

### Plasmid DNA, molecular biology and virus production

A CRISPRv2 construct bearing sgRNA targeting genes of interest was generated as follows. 5 µg of CRISPRv2 was digested with BsmBI at 37 °C for 30 min. The digested vector was gel-purified and eluted in pure water. Each pair of sgDNA oligos were phosphorylated and annealed by incubating 1µl of sgRNA oligo 1 or 2 (100 µM) with 1µl 10X T4 ligation buffer in the presence of 0.5µl T4 polynucleotide kinase in 10µl. The reaction was placed in a thermocycler using the following parameters: 37 °C for 30 min, 95 °C for 5 min, and ramped down to room temperature at 5 °C/min. The annealed sgDNA oligo was then ligated with the BsmBI-digested CRISPRv2 to create CRISPRv2-sgDNA constructs targeting *SLC35B2* and *MYO7B*. KO lentivirus were produced by transfecting 1 million HEK293FT cells in a 3.5 cm dish with 0.4 µg pVSV-G, 0.6 µg psPAX2 and 0.8 µg CRISPRv2-sgRNA. Transfected cells were incubated with 3 ml fresh DMEM medium for 72 h before virus were harvested. shRNA knockdown virus was produced in a similar manner except that a pair of shRNA-coding oligos were annealed and ligated with Age1-EcoR1digested pLKO.1 construct. SLC35B2 rescue plasmid was generated by PCR-amplifying the SLC35B2 cDNA sequence with a FLAG tag-coding sequence and a NheI site appended at the 5’-end and an EcoRI site at the 3’-end. The PCR fragment was digested with NheI and EcoRI and further purified. The purified DNA fragment was then inserted into pLJM1. Five non-sense mutations were introduced at the sgRNA targeting sequence by site directed mutagenesis to avoid Cas9 recognition of the rescue DNA. PCR mutagenesis was performed using QuickChange II site directed mutagenesis kit (Agilent) according to the manufacturer’s protocol.

### Protein purification, labeling, and PFF production

Human α-Synuclein (α-Syn) protein was expressed in BL21(DE3)RIL competent *E. coli* and purified following an established protocol (27). Briefly, 1L *E. coli* culture was induced to express α-Syn with IPTG 0.5mM at 20 °C overnight. Cells were pelleted down at 5000 rpm for 20 min then resuspended with high salt buffer (750 mM NaCl, 10 mM Tris, pH 7.6, 1 mM EDTA) with protease inhibitors and 1 mM PMSF (50 mL for 1L of culture). Resuspended cells were sonicated with 0.25 inch probe tip at 60% power for a total time of 10 min. Sonicated cell lysate was boiled for 15 min to precipitate unwanted proteins and cooled on ice for 20 min. The lysate was then centrifuged at 6,000 x g for 20 min. The supernatant containing α-Syn was collected and dialyzed in TNE buffer (10 mM Tris, pH 7.6, 25 mM NaCl, 1mM EDTA). Isolated proteins were further fractionated with a BioRad NGS chromatography system first through a size exclusion column in TNE buffer, followed by an ion exchange Mono-Q column. Protein was eluted in TNE buffer containing increasing concentrations of NaCl (25mM-1M). Purified monomeric α-Syn was dialyzed extensively in the phosphate buffered saline (PBS) followed by labeling with 3-fold molar of pH-Rodo Red or Alexa596 succinimidyl ester fluorescence dye (Thermo Fisher) for 4 h at room temperature. Unconjugated dye was removed by dialysis in PBS. The labeled α-Syn was adjusted to 5 mg/ml in 500 µl volume and was placed in a thermomixer shaker at 1,000 RPM at 37 °C for 7 days to generate protein fibrils. Preformed α-synuclein fibrils (α-Syn PFF) were stained with 3% uranyl acetate and examined by EM to verify the formation of α-Syn PFF. α-Syn PFF were stored in a −80 °C freezer in small aliquots. GPF+ was expressed and purified according to a previously published protocol (34).

### α-Synuclein PFF uptake assay and drug treatment

α-Syn PFF was sonicated right before each use as previously reported (27). For FACS analysis, cells were incubated with 800 nM α-Syn PFF for 3h whereas in imaging experiments, cells were incubated with 200-400 nM α-Syn PFF for 3h in regular DMEM medium. To distinguish between internalized α-Syn PFF or cell surface bound PFF, cells were washed with PBS containing 4 µM Heparin prior to imaging experiments as described in the figure legends. To examine the effect of drug treatment on α-Syn PFF uptake, cells were incubated with drug 30min prior to the addition of α-Syn PFF. The following drugs were used, Dynasore (80 µM), Dynole 34-2 (15-25 µM), Jasplakinolide (100 nM), Latrunculin A1 (1 µM) and methyl-β-cyclodextrin (5 mM).

α-Syn PFF uptake in primary neurons was performed as follows. Primary hippocampal neuron cultures were prepared from P0-1 murine pups. Hippocampi were dissected in Hanks’ balanced salt solution (HBSS) and washed with MEM (Gibco) twice. The hippocampi were incubated with 0.25% trypsin (Gibco) containing 100 µg/mL DNase-I (Sigma) for 10 min at 37 °C. Trypsin was then inactivated by addition of MEM containing 10% FBS. After washing with MEM three times, the hippocampal cells were dissociated by pipetting several times in MEM containing 100 µg/mL DNase-I. Cells were then centrifuged at 300 g for 5 min and resuspended in the plating medium (MEM containing 10% FBS, 1 mM sodium pyruvate (Sigma), 2 mM L-glutamine (Sigma), 100 µg/mL Primocin (Invitrogen) and 0.6% glucose). The cell solution was passed through a 70 µm strainer (VWR) once to filter out any cell clumps. Cells were seeded into 24 well plates pre-coated with poly-D-lysine (50 µg/ml) and laminin (2 µg/mL). Generally, we seed all hippocampal cells obtained from 4 pups to prepare one 24-well plate culture. After incubation at 37 °C with 5% CO2 for 24 h, the culture medium was changed to the neuronal culture medium (Neurobasal media (Invitrogen) containing 2% B27 (Thermo Fisher), 0.5 mM L-glutamine and 100 µg/mL Primocin) to support the growth of hippocampal neurons. For α-Syn PFF treatment, the medium at DIV7-9 was replaced with fresh neuronal culture medium containing α-Syn PFF (400nM). Cells were stained with DAPI and imaged by a confocal microscopy. To perform shRNA-mediated gene silencing, neurons were seeded in 12-well plate and infected with 500 µl shRNA-expressing lentivirus at day-in-vitro 3 (DIV3). Cells were placed in an incubator for an additional 5 days before examining the silencing effect.

### CRISPR knockout screen

The GeCKO V2 human library A and B containing 112417 sgRNAs targeting 19052 genes as well as 1000 nontargeting control sgRNAs was obtained from Addgene (Cat.# 1000000048 and 1000000049). Plasmid DNA was amplified and lentivirus was produced and amplified using HEK293FT cells following the provided protocol (51). The virus titer (or multiplicity of infection. M.O.I.) was determined as described previously (51).

To generate cells expressing the KO library, 60 million HEK293293T cells were resuspended at 3X10^6^ cells/ml in fresh DMEM medium containing polybrene (8 µg/ml). Cells were seeded into 12-well plates at 1ml/well for a total of 20 wells. A virus-medium mixture was prepared by addition of 300 µl virus solution into 10 ml fresh DMEM medium containing polybrene. 500 ml virus-medium mixture was added to each well. Cells were spin-infected at 2,000 rpm at 37 °C for 2 hours, then incubated in cell culture incubator for 1 h before changing the medium. After 24 hours, cells were pooled and incubated with DMEM containing puromycin (0.35 µg/ml) for 96 hours to generate cells stably expressing the sgRNA KO library. To screen for mutant cells defective in α-Syn PFF uptake, 60 million library cells were treated with 500nM α-Syn PFF pH-Rodo for 3 hours before FACS analysis. WT HEK293T cells treated with α-Syn PFF in parallel were used as a reference for cells with normal uptake. Cells with lower fluorescence intensity compared to WT cells were identified and sorted individually into 96 wells by BD LSR-II flow cytometry (NHLBI flow cytometry core). When cell clones were formed, they were treated with 500nM α-Syn PFF pH-Rodo 3h again and analyzed with a flow cytometer to identify clones truly defective in α-Syn PFF uptake. Genomic DNA was extracted from cells by genomic purification kit (Thermo Fisher) and a 300 bp DNA fragment containing sgRNA sequence was amplified PCR, cloned into pCR4-TOPO TA vector and sequenced.

### qRT-PCR analysis of knockout or knockdown cells

Total RNA was extracted from 3 million HEK293T cells or 0.75 million primary neurons using TriPure reagent (Roche) and purified using RNeasy MinElute Cleanup Kit (Promega) following standard protocols. The RNA concentration was measured by Nanodrop 2000 UV spectrophotometer and 1µg total RNA was converted to cDNA using the iScript Reverse Transcription Supermix (BioRad) system. 1 µL cDNA was used to perform qPCR using SsoAdvanced SYBR Green supermix kit (BioRad) on a CFX96 machine (BioRad). Data were analyzed using BioRad CFX manager 3.0 software. GADPH was used as a reference gene for quantification of gene expression levels. Primers used to generate KO cells and for qRT-PCR were listed in SI Table 2:

### Structural modeling

The models of α-Syn PFF in complex with heparin were obtained using the open source program ClusPro protein-protein docking platform (http://cluspro.org/login.php) (52–54). Previously published α-Syn fibril structures (PDB: 6CU7 6CU8) were used as the model protein and heparin was set as the ligand (35). The analyses resulted in 5 and 9 clusters for “rod” and “twisted conformer”, respectively. These clusters were further divided into 3 groups, representing 3 modes of interactions as shown by models in Figure 4A-C, Supplementary SI Fig. S3A, and SI Fig. S3B, respectively. Clusters within the same group are similar to each other as they all use the same site formed by either one or two K-T-K motifs to bind heparin, which adopts slightly different orientations within the same groups. We chose to present the clusters with the heparin in parallel to the binding groove. For analysis of hydrogen bonding potential, we used LigPlot program (55).

### HSPGs staining, and α-Syn PFF binding to cell surface and GFP+ binding to cell surface

Cells were fixed with 4% paraformaldehyde in PBS at room temperature for 15 min, washed extensively with PBS, and incubated in PBS containing 5% fetal bovine serum for 10min. A mouse monoclonal antibody to HSPGs (US Biological) was used to stain the cell surface at 1/100 dilution for 4 h. Cells were then stained with a secondary antibody conjugated with Alexa _596_ (Thermo Fisher) at 1: 5,000 dilution for 1 h at room temperature.

To examine α-Syn PFF binding to the cell surface, cells were incubated with 400nM α-Syn PFF-Alexa_594_ on ice for 30 min. Cells were then washed with ice-cold PBS 3 times to remove unbound α-Syn PFF and fixed with PBS containing 4% paraformaldehyde at room temperature for 15min. Cell nuclei were stained with Hoechest 33342 (Invitrogen) in PBS at 1µg/ml concentration prior to imaging. To examine the inhibitory effect of heparin on α-Syn PFF binding, cells were pre-treated with 4 µM heparin (Sigma) before adding α-Syn PFF.

For live-cell imaging experiments, cells were seeded in 8-well ibidi chamber at 5,000 per well. At 24 h post seeding, cells were treated with pHrodo- or Alexa594-labeled α-Syn PFF or with GFP+. The medium was then replaced with a DMEM-based imaging medium lacking phenol red. Live-cell images were acquired using a Zeiss LSM780 laser scanning confocal microscope (63X/1.46 objective) with the Zen program (Zeiss).

### TIRF imaging of fluorescence-tagged clathrin light chain and MYO7B

TIRF imaging was performed on a Zeiss microscope equipped with a 63X/1.46 TIRF objective lens. The TIRF microscope was calibrated using fluorescent beads provided by Zeiss. The critical angles were determined according to the manufacturer’s manual (Zeiss). Cells stably expressing fluorescently tagged-CLC or MYO7B were seeded in a 4-well ibidi chamber (refractive index 1.5) at 5000 cells per well and incubated overnight before imaging.

### Electron microscopy analysis of α-Syn PFF and CCPs

5mg/ml α-Syn PFF was diluted 10-fold in PBS and loaded onto an EM grid. The grid was then washed with pure water and stained with 3% uranyl acetate for 1 min before imaging processing with a Morgagni 268 transmission electron microscope.

For transmission EM analysis of CCPs, cells were grown on fibronectin-coated coverslips and fixed in 2.5% glutaraldehyde, 2% formaldehyde, 0.1 M cacodylate, 2 mM calcium chloride pH 7.4 (CacCl) for 15 min at room temperature followed by 45 min on ice. Coverslips were washed 3 times for 5 min each with CacCl, treated with 1% tannic acid for 30’, washed and post-fixed for 30’ in 0.5% osmium tetroxide/0.5% potassium ferrocyanide in the same buffer. Cells were washed twice with CacCl, twice with 50 mM sodium acetate pH 5.2 and stained for 30 min with 2% uranyl acetate. After washing 5 times with water, the samples were dehydrated through a series of increasing concentration of ethanol (30%, 50%, 70%, 90%, 3 times 100% anhydrous) and embedded in EMBed 812 epoxy resin (EMS, Hatfield, PA). After resin polymerization, the coverslip was removed by hydrofluoric acid, sample blocks cut out and mounted on a holder. Ultrathin sections (70-80 nm thick) were cut parallel to the plane of the coverslip and mounted on formvar/carbon-coated EM grids. Sections were stained with lead citrate and imaged in an FEI Tecnai 20 transmission electron microscope operated at 120 kV. Images were recorded on an AMT XR81 wide-field CCD camera.

### Image processing and statistical analysis

Images were processed using the Zeiss Zen software. To measure fluorescence intensity, we used the Fiji software. Images were converted to individual channels and region of interests were drawn for measurement. To measure the size of CLC pits, particles were identified by intensity threshold. Particles above the cutoff value were identified and measured. To measure the plasma membrane dynamics, the Nikon Element software was used. For each cell, at least 10 GFP+-stained membrane dots were identified and tracked by the software, and the averaged velocity was calculated. Statistical analyses were performed using either Excel or Prism 8. Values are the mean ± SEM, calculated by GraphPad Prism 8, and *p*-values were calculated by the GraphPad Prism 8 or by Student’s t-test using Excel. Images were prepared with Adobe Photoshop and graphs were plotted by GraphPad Prism 8.

## Data Availability

All data, associated protocols, methods, and sources of materials can be accessed in the text or *SI Appendix*.

## Acknowledgements

We thank Dr. R. Tycko and Dr. U. Ghosh (NIDDK) for assistance in protein negative stain, J. Reece at the NIDDK imaging core for imaging assistance, D. Lu for assistance with data processing, H. Meyer (University of Duisburg Essen) for critical reading of the manuscript, M. Tyska (Vanderbilt University) for MYO7B WT and mutant cDNAs. Q. Zhang, Y. Xu, J. Lee, and Y. Ye are supported by the intramural research program of NIDDK (DK075143), X. Wu by the intramural research program of NHLBI, M. Jarnik and J. S. Bonifacino by the intramural research program of NICHD (ZIA HD001607) in NIH, J. Shen by NIH grants AG061829 and GM126960.

## Author contributions

Q. Zhang, Y. Xu, and Y. Ye designed the research, performed experiments and analyzed the data. M. Jarnik and J.S. Bonifacino performed EM analyses of CCPs, X. Wu performed super-resolution microscopy, J. Shen confirmed CME of α-Syn PFF and assisted in CRISPR screen. Q. Zhang and Y. Ye wrote the paper and all authors participated in editing the manuscript.

## Competing interests

The authors declare no competing financial interest.

