## Supplemental figures for "A Myosin-7B dependent endocytosis pathway mediates cellular entry of α-Synuclein fibrils and polycation-bearing cargos"

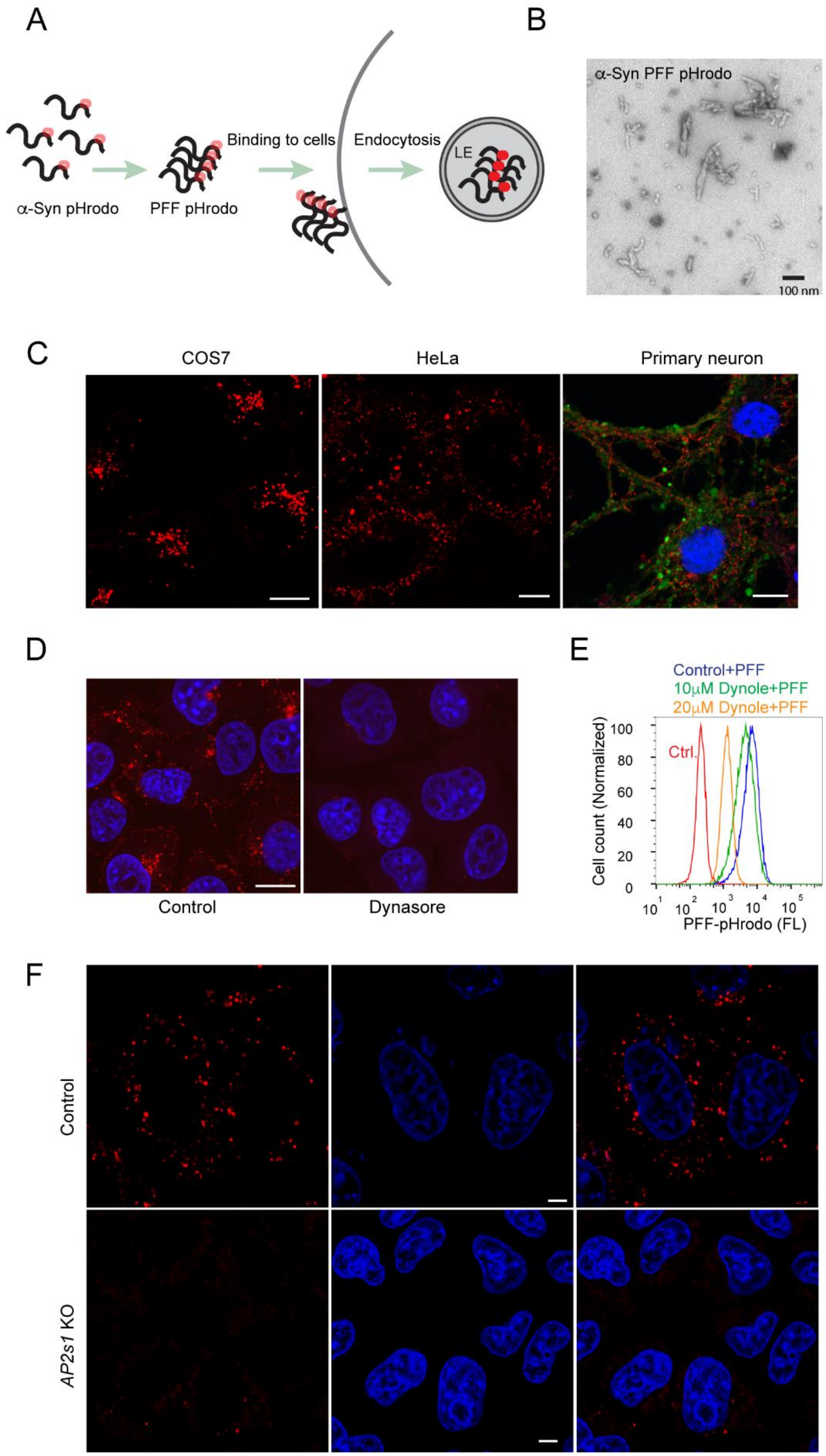
A
α-Syn pHrodo
PFF pHrodo
Binding to cells
Endocytosis
LE
B
α-Syn PFF pHrodo
100 nm
C
COS7
HeLa
Primary neuron
D
Control
Dynasore
E
Control+PFF
10μM Dynole+PFF
20μM Dynole+PFF
Ctrl.
Cell count (Normalized)
PFF-pHrodo (FL)
F
Control
AP2s1 KO

**Fig. S1. CME mediates α-Syn PFF uptake.**
**(A)** An illustration of the strategy for tracking α-Syn PFF endocytosis. LE, late endosome
**(B)** EM analysis of sonicated α-Syn PFF.
**(C)** α-Syn PFF pHrodo is efficiently internalized by various cell types including COS-7, HeLa and primary neurons. For primary neurons, cells were infected with lentivirus expressing GFP under the neuronal specific *SYN1*-promoter and stained with DAPI to reveal the nuclei (blue). Scale bars, 10µm.
**(D-F)** α-Syn PFF endocytosis is largely mediated by CME. **(D)** COS-7 cells pre-treated with DMSO as a control or with the dynamin inhibitor Dynasore (80µM, 1 h) were incubated with 400nM α-Syn PFF $Alexa_{594}$ for 3h. Cells were washed with heparin and stained with DAPI before imaging. Scale bar, 10µm. **(E)** HEK293T cells pre-incubated with DMSO or the indicated concentration of the dynamin inhibitor Dynole 34-2 (Dynole) were incubated with α-Syn PFF pHrodo and analyzed by FACS. **(F)** Control or *AP2S1*-KO HeLa cells were incubated with pHrodo-labeled α-Syn PFF (red) and then stained with DAPI (blue) prior to fluorescence imaging. Scale bar, 5µM.

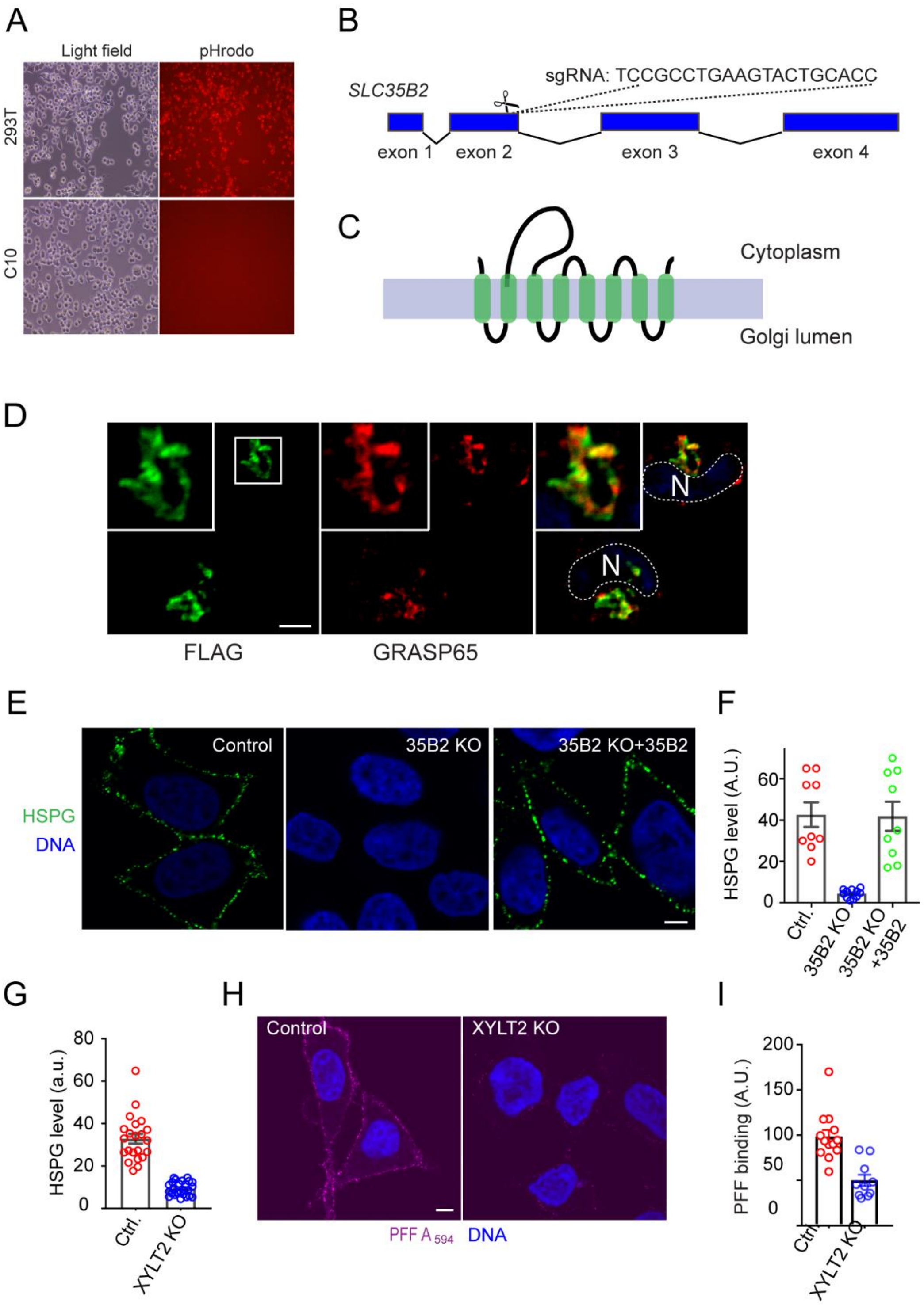
A
Light field
pHrodo
293T
C10
B
SLC35B2
sgRNA: TCCGCCTGAAGTACTGCACC
exon 1
exon 2
exon 3
exon 4
C
Cytoplasm
Golgi lumen
D
N
N
FLAG
GRASP65
E
Control
35B2 KO
35B2 KO+35B2
HSPG
DNA
F
HSPG level (A.U.)
60
40
20
0
Ctrl.
35B2 KO
35B2 KO +35B2
G
HSPG level (a.u.)
80
60
40
20
0
Ctrl.
XYLT2 KO
H
Control
XYLT2 KO
PFF A 594
DNA
I
PFF binding (A.U.)
200
150
100
50
0
Ctrl.
XYLT2 KO

**Fig. S2. HSPG mediates α-Syn PFF uptake.**
**(A)** C10 cells are defective in the uptake of pHrodo-labeled α-Syn PFF. Unless otherwise specified in the figure legends, cells were incubated with α-Syn PFF at 400nM for 3h. Live-cells were imaged by an epi-fluorescence microscope.
**(B)** Mapping the identified SLC35B2 sgRNA.
**(C)** The predicted membrane topology of SLC35B2.
**(D)** SLC35B2 is a Golgi-localized protein. *SLC35B2*-KO cells expressing FLAG-SLC35B2 were stained with antibodies to FLAG epitope and to the Golgi protein GRASP65. The insets show an enlarged view of the boxed area. N, nuclei. Scale bar, 5µm.
**(E, F)** SLC35B2 is required for HSPG biosynthesis. Control or *SLC35B2*-KO cells with or without reconstitution of SLC35B2 were stained with antibody to HSPG (Green) and with DAPI (blue). Scale bar, 5µm. The graph in **(F)** shows the relative HSPG levels in randomly selected cells from two experiments. Error bar, SEM.
**(G-I)** XYLT2 is required for α-Syn PFF endocytosis. The graph in **(G)** quantifies the HSPG levels from confocal images of control and *XYLT2*-KO cells stained with the HSPG antibody. Error bar, S.E.M**, (H)** Control or *XYLT2*-KO cells were incubated with 200nM α-Syn PFF $Alexa_{594}$ for 30 min before imaging. The graph in **(I)** shows the relative α-Syn PFF level on the cell surface from two experiments. Error bar, SEM.

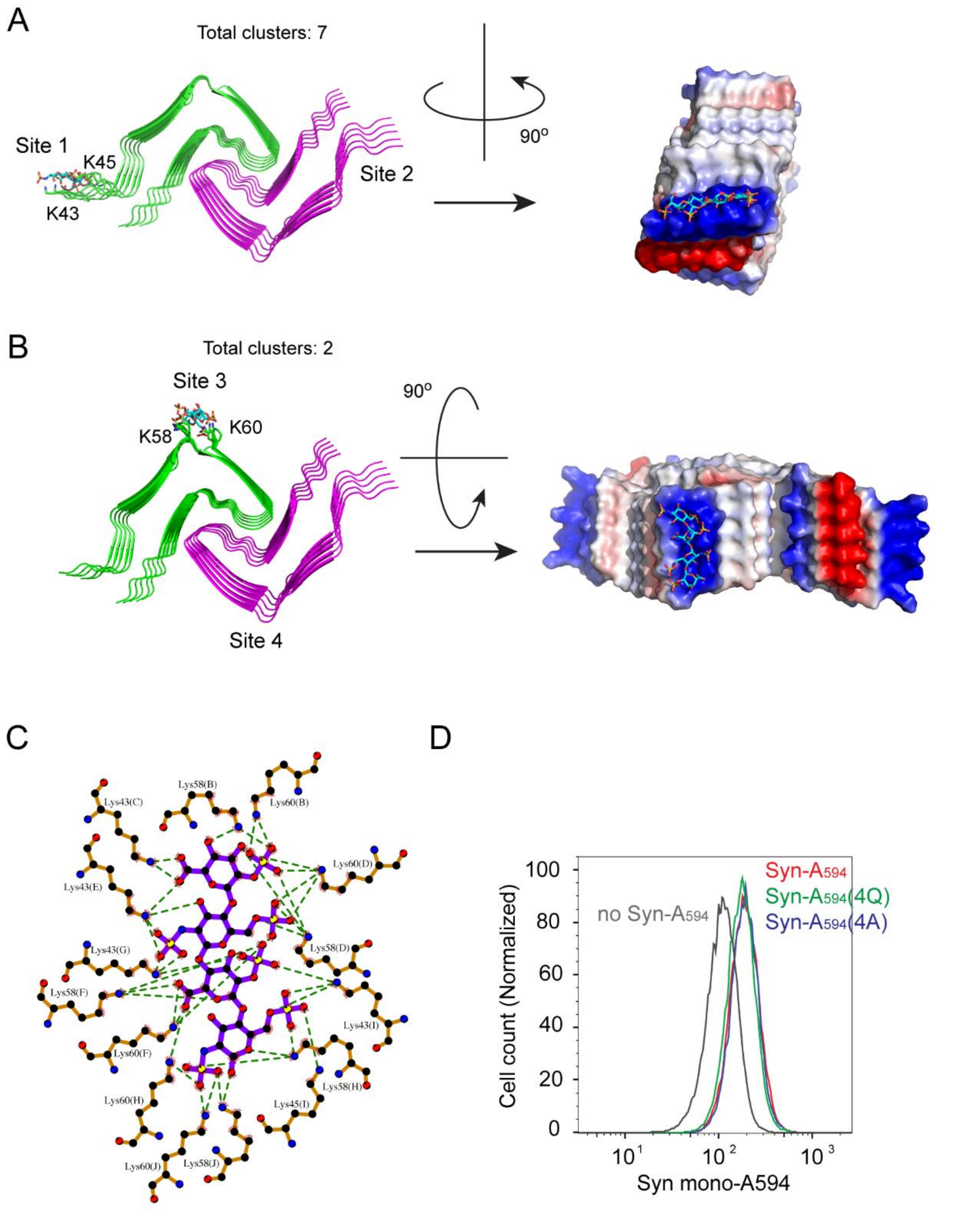


**Fig. S3. α-Syn PFF interacts with heparin using two K-T-K motifs.**
(A, B) In the twisted conformer, each dimeric fibril can have four α-Syn PFF binding sites, but each site uses only one K-T-K motif to interact with heparin and thus is expected to have lower affinity compared to the binding site formed by two K-T-K motifs in the rod conformer.
**(C)** Potential electrostatic and hydrogen bond interactions between heparin and the rod conformer of α-Syn PFF as calculated by LigPlot (1).
**(D)** The K-T-K motifs are not required for endocytosis of monomeric α-Syn. Endocytosis of $Alexa_{594}$-labeled WT or mutant α-Syn monomers (4A and 4Q, 200nM each) was measured after overnight uptake by FACS.

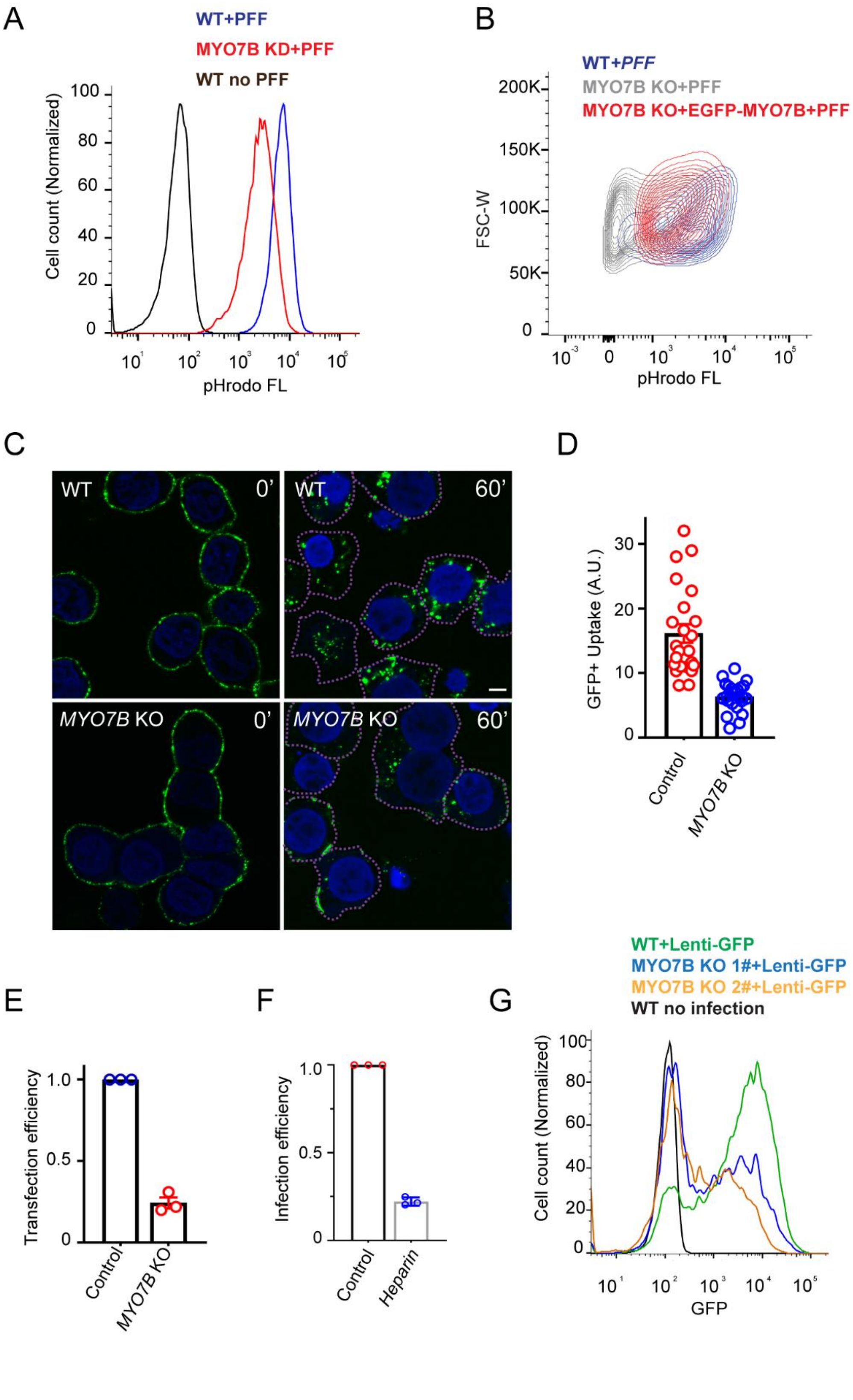
A
WT+PFF
MYO7B KD+PFF
WT no PFF
Cell count (Normalized)
pHrodo FL
B
WT+PFF
MYO7B KO+PFF
MYO7B KO+EGFP-MYO7B+PFF
FSC-W
pHrodo FL
C
WT 0'
WT 60'
MYO7B KO 0'
MYO7B KO 60'
D
GFP+ Uptake (A.U.)
Control
MYO7B KO
E
Transfection efficiency
Control
MYO7B KO
F
Infection efficiency
Control
Heparin
G
WT+Lenti-GFP
MYO7B KO 1#+Lenti-GFP
MYO7B KO 2#+Lenti-GFP
WT no infection
Cell count (Normalized)
GFP

**Fig. S4. MYO7B facilitates endocytosis of α-Syn PFF and polycation-bearing cargos.**
**(A)** α-Syn PFF uptake was monitored in WT or *MYO7B*-knockdown (KD) HEK293T cells by FACS. Cells were incubated with pHrodo-labeled α-Syn PFF (400nM, 4 h). WT untreated cells were used as a negative control.
**(B)** Re-expression of EGFP-MYO7B in *MYO7B*-KO cells rescues α-Syn PFF uptake. Shown are FACS contour maps of *WT* (blue), *MYO7B*-KO (grey), or *MYO7B*-KO HEK293T cells re-expressing low levels of EGFP-MYO7B (red) after treatment with pHrodo α-Syn PFF (400nM, 4 h).
**(C, D)** *MYO7B*-KO cells are defective in uptake of GFP+. **(C)** Cells incubated with GFP+ (200nM) for 5min were imaged directly (t=0') or chased at 37 °C for 60 min (t=60') followed by wash with heparin to remove cell surface-bound GFP+. Cells were stained with DAPI (blue) and imaged by confocal microscopy. The graph in **(D)** quantified the level of GFP+ in individual cells. Error bars, SEM.
**(E)** *MYO7B*-KO cells are defective in DNA transfection. Cells were transfected with a mCherry plasmid for 24 h and analyzed by FACS. error bars, SEM, n=3.
**(F, G)** MYO7B is also required for HSPG-dependent lentiviral infection. **(F)** Cells pre-treated with heparin or untreated were infected with lentiviral particles bearing an EGFP reporter. The EGFP positive cells were analyzed by FACS 24 h post-infection. **(G)** WT or two clones of *MYO7B*-KO cells were infected with EGFP-bearing lentiviral particles and analyzed by FACS for GFP expression. Uninfected WT cells were used as a negative control.

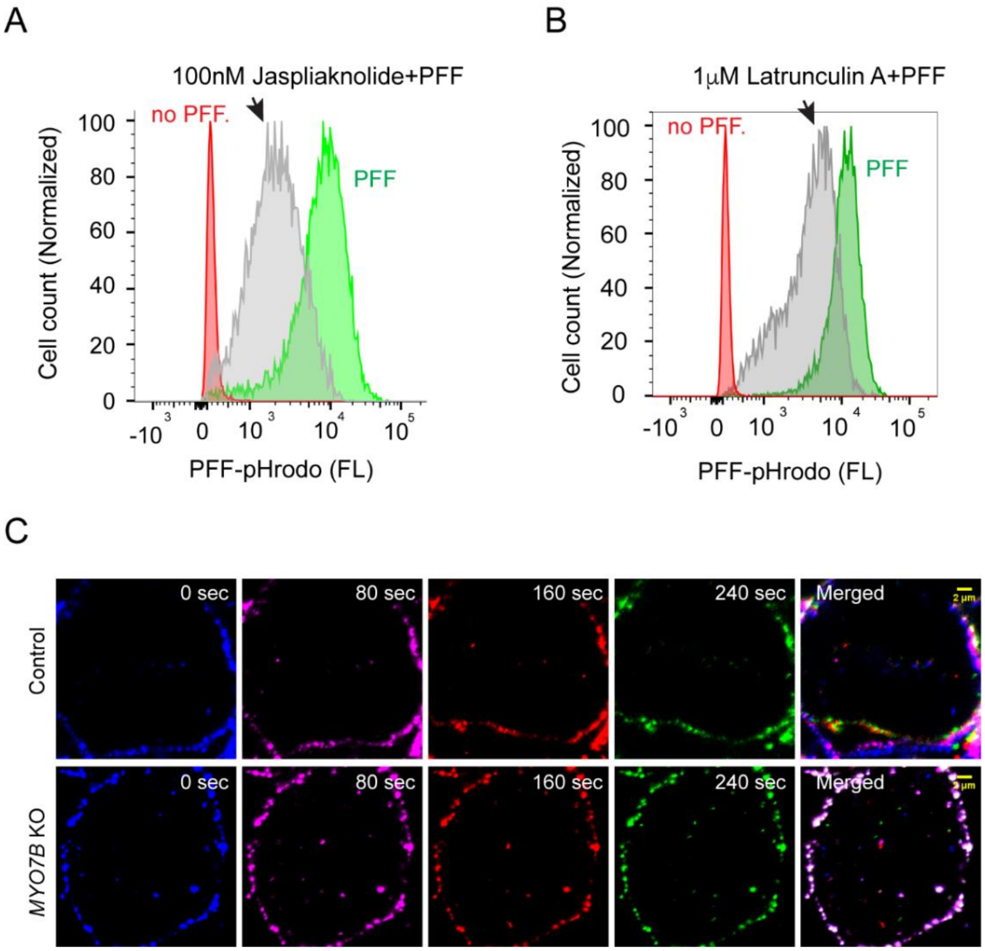


**Fig. S5 Actin-dependent membrane dynamics is required for α-Syn PFF endocytosis.**
**(A, B)** Actin-binding compounds inhibit α-Syn PFF endocytosis. HEK293T cells were treated with the indicated actin-binding compounds (arrows) or left untreated (green) for 30min before being incubated with pHrodo α-Syn PFF for 3h. Cells that have not been exposed to α-Syn PFF were used as a negative control (red).
**(B)** MYO7B inactivation affects plasma membrane dynamics. Control or *MYO7B*-KO cells were stained with GFP+ and imaged by time-lapse confocal microscopy. Images at the indicated time frames of a video were pseudo-colored and merged to reveal the movement of plasma membrane. Scale bars, 2µm.

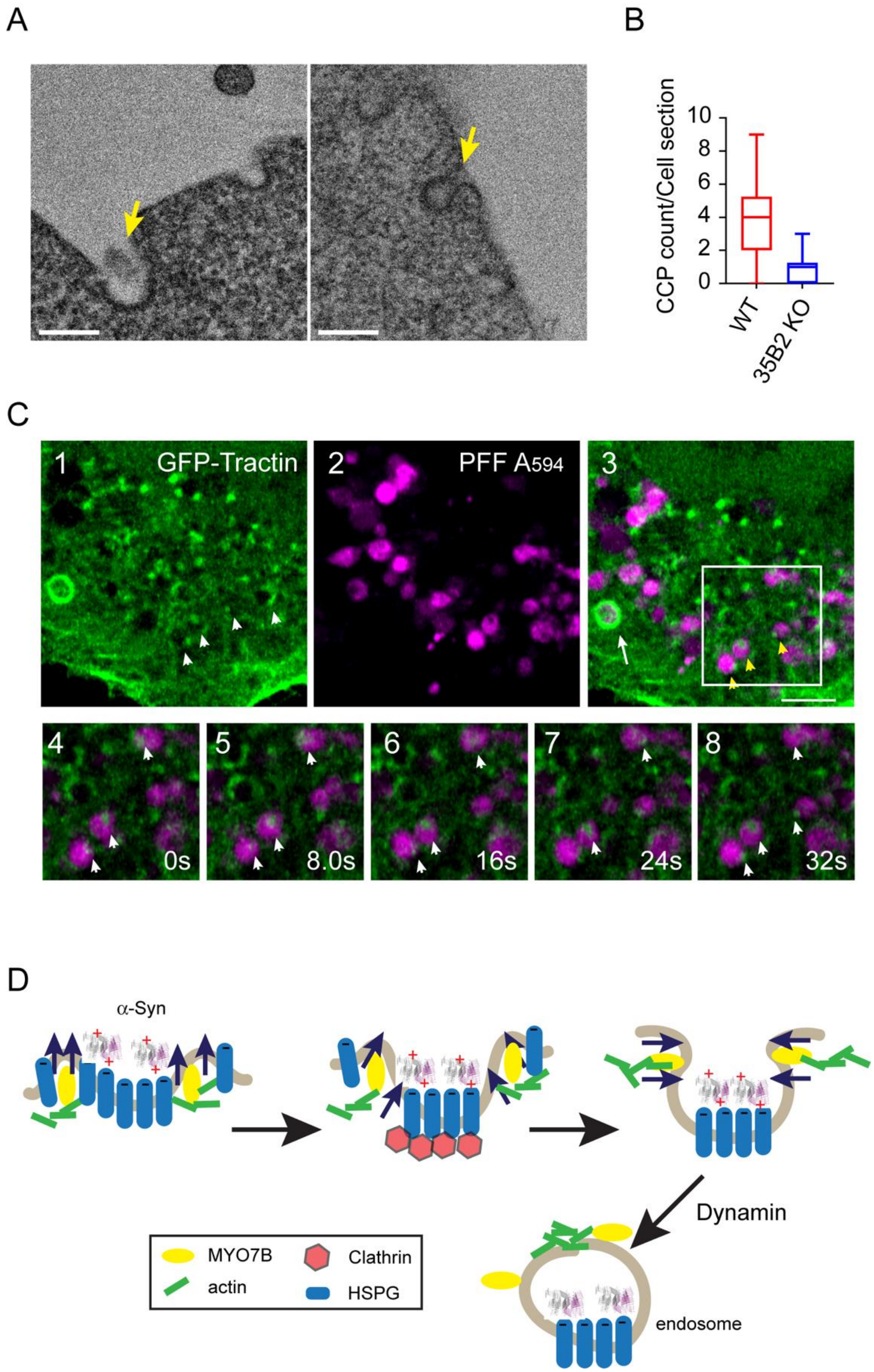

A
B
CCP count/Cell section
10
8
6
4
2
0
WT
35B2 KO
C
1
GFP-Tractin
2
PFF A594
3
4
5
6
7
8
0s
8.0s
16s
24s
32s
D
α-Syn
Dynamin
MYO7B
Clathrin
actin
HSPG
endosome

**Fig. S6**. **A working model for endocytosis of α-Syn PFF and other HSPG cargos.**
**(A)** Representative EM images of CCPs from WT and *SLC35B2KO* cells. Cells were treated with 400 nm α-Syn PFF 3h before EM analysis. Arrows indicate CCPs. Scale bars, 200nm.
**(B)** *SLC35B2*-KO cells have a reduced number of CCPs. The box-whisker plot shows the relative number of CCPs per cell section**.** n=14 and 30 for WT and *SLC35B2*-KO cells, respectively.
**(C)** Actin filaments are detected on α-Syn PFF bearing vesicles. Cells transiently expressing GFP-Tractin (green) were incubated with α-Syn PFF $Alexa_{594}$ (200nM, 3h) (magenta) and then imaged by an Airyscan confocal microscope. Lower panels show frames from a time-lapse video, focusing on the boxed area in panel 3. Scale bar, 5µm.
**(D)** α-Syn PFF binding triggers HSPGs oligomerization. MYO7B assembles actin filaments at the edge of HSPG oligomers, pushing the plasma membrane outwards while HSPGs recruit clathrin (possibly via an adaptor not shown). Continued assembly of actin filaments eventually drives the membrane around α-Syn PFF, which leads to dynamin-mediated membrane severance.

**SI Table 1 A list of reagents**

| REAGENTS | SOURCE |
|---|---|
| **PLASMIDS** | |
| **pEGFP-MY07B FL** | A gift from Matthew Tyska(2) |
| **pEGFP-MY07B FL (N207A)** | A gift from Matthew Tyska(2) |
| **pEGFP-MY07B FL (I482A)** | A gift from Matthew Tyska(2) |
| **pEGFP-MY07B FL (E442A)** | A gift from Matthew Tyska(2) |
| **pEGFP-MY07B C TAIL** | A gift from Matthew Tyska(2) |
| **pEGFP-MY07B C TAIL (890STOP)** | This study |
| **FUGW** | A gift from David Baltimore Addgene (#14883)(3) |
| **pCMV-VSV-G** | A gift from Bob Weinberg Addgene (#8454)(4) |
| **pSPAX2** | A gift from Didier Trono Addgene (#12260) |
| **pEGFP-C1** | Clonetech |
| **LENTICRISPR V2** | A gift from Feng Zhang Addgene (#52961)(5) |
| **pEGFP-C-SLC35B2** | This study |
| **pEGFP-N-SLC35B2** | This study |
| **pEGFP-C1 F-TRACTIN** | A gift from Dyche Mullins Addgene (#58473)(6) |
| **GFP-FKBP-CLATHRIN LIGHT CHAIN** | A gift from Stephen Royle Addgene(#59353)(7) |
| **mCHERRY-CLATHRIN LIGHT CHAIN** | A gift from Michael Davidson Addgene (#55019)(8) |
| **pEGFP-N1 DYNAMIN1** | A gift from Justin Taraska Addgene (#120313)(9) |
| **DYN2-PMCHERRYN1** | A gift from Christien Merrifield Addgene (#27689)(10) |
| **PLJM1-EGFP** | A gift from David Sabatini Addgene(#19319)(11) |
| **mCHERRY-N1-TRACTIN** | This study |
| **PLJM1-FLAG-SLC35B2-RESCUE** | This study |
| **PLKO.1** | A gift from David Root Addgene(#10878)(12) |
| **CHEMICALS** | |
| **PUROMYCIN** | Sigma |
| **BAFILOMYCIN A1** | Millipore |
| **PH RODO-RED SUCCINIMIDYL ESTER** | ThermoFisher Scientific |

| | |
|---|---|
| **ALEXA 596 SUCCINIMIDYL ESTER** | ThermoFisher Scientific |
| **DYNOLE 34-2** | TOCRIS |
| **DYNASORE** | TOCRIS |
| **JASPLAKINOLIDE** | TOCRIS |
| **LATRUNCULIN A** | TOCRIS |
| **HOECHEST 33342** | ThermoFisher Scientific |
| **METHYL-β-CYCLODEXTRIN** | Sigma |
| **IPTG** | TOCRIS |
| **ANTIBODIES** | |
| **H1890 MOUSE ANTI-HEPARAN SULFATE (10E4 EPITOPE) ANTIBODY** | US Biological |
| **Alpha-Synuclein ANTIBODY (211)** | Santa Cruz Biotechnology |

**SI Table 2 A list of primers used in the study**

| | |
|---|---|
| MYO7B KO qPCR F: | 5’-TATGGGCGAGCTGCCCCCGC-3’ |
| MYO7B KO qPCR R: | 5’-GCGCGCGCCCTCGATCACCC-3’ |
| USH1C KO qPCR F: | 5’-ACATCGGCCTGATCCCCGTGAAA-3’ |
| USH1C KO qPCR R: | 5’-AATCTGGTCCCCTATCTCCACTC-3’ |
| XYLT2 KO qPCR F: | 5’-TGGTGGATGGCGGTTCTGAC-3’ |
| XYLT2 KO qPCR R: | 5’-GCTTGAAGTCGTTGGGGGAG-3’ |
| SLC35B2 KO qPCR F: | 5’-ACCTCCTGGTGCAGTACTTC-3’ |
| SLC35B2 KO qPCR R: | 5’-ACAGGCTGGCAAAGGAGTAC-3’ |
| mMYO7B KD qPCR F: | 5’-GAAAGCGGCTGAGCAGAATG-3’ |
| mMYO7B KD qPCR R: | 5’-CACCTGATAGGGAAGTGTCA-3’ |
| mSLC35B2 KD qPCR F: | 5’-CCTTTGCCAGTCTGTCAAAT-3’ |
| mSLC35B2 KD qPCR R: | 5’-ATAGGCAAACAGGGCATCCT-3’ |
| SLC35B2 sgRNA : | 5’-TCCGCCTGAAGTACTGCACC-3’ |
| MYO7B sgRNA: | 5’-GCGGGGGCAGCTCGCCCATA-3’ |
| XYLT2 sgRNA: | 5’-TGGTGGATGGCGGTTCTGAC-3’ |
| USH1C sgRNA: | 5’-TCGGCCTGATCCCCGTGAAA-3’ |
| MYO7B siRNA1: | 5’-AGAGCAUCCUUCUAGCCUATT-3’ |
| MYO7B siRNA2: | 5’-CAGACACCAUACUCCAUUATT-3’ |
| MYO7B siRNA3: | 5’-GCCCAGAAGUUUAUAGACATT-3’ |

**Movie S1 (separate file).** A time-lapse TIRF video shows the dynamic interaction of MYO7B with the plasma membrane in a U2OS cell stably expressing EGFP-WT MYO7B.
**Movie S2 (separate file).** A time-lapse TIRF video shows the plasma membrane-localized MYO7B CT in a U2OS cell stably expressing EGFP-CT MYO7B.
**Movie S3 (separate file).** A time-lapse fluorescence confocal video shows a WT HEK293T cell stained with GFP+. The video was processed by Nikon element to track mobility of GFP+-labeled plasma membrane puncta.
**Movie S4 (separate file).** A time-lapse fluorescence confocal video shows a *MYO7B*-KO HEK293T cell stained with GFP+. The video was processed by Nikon element to track mobility of GFP+-labeled plasma membrane puncta. Note that the plasma membrane is almost completely static.
